# Conserved control of cell differentiation across divergent brown algal body plans revealed by diploid genome editing

**DOI:** 10.64898/2026.11.25.769193

**Authors:** Anagha Kerur, Morgane Raphalen, Susana M. Coelho, Kenny A. Bogaert

**Affiliations:** Department of Algal Development and Evolution, Max Planck Institute for Biology, Max-Planck-Ring 5, 72076 Tübingen, Germany; Marine Biological Station Carolinensiel, Institute of Integrative Cell Biology and Physiology, University of Münster, Friedrichsgroden 16, 26409 Wittmund, Germany

**Keywords:** CRISPR, brown algae, cell differentiation, reverse genetics, TBCC, *Dictyota*

## Abstract

Brown algae evolved complex multicellularity independently of animals, plants and fungi, yet whether developmental mechanisms are conserved across their deeply divergent and morphologically diverse lineages remains poorly understood. In the filamentous brown alga *Ectocarpus*, the evolutionarily conserved protein DISTAG (DIS) is required for basal cell identity, but its function in more complex body plans is unknown. Here, we investigate the role of DISTAG in *Dictyota*, a distantly related brown alga that develops a complex parenchymatous thallus. To enable functional genetic analysis in this species, we developed a DNA-free CRISPR–Cas genome-editing approach that exploits the short window between gamete release and cell wall formation to introduce Cas ribonucleoproteins into zygotes. This strategy allows edited alleles to be maintained in heterozygous sporophytes and recovered in haploid progeny following meiosis, enabling genetic analysis across generations. Using this approach, we show that loss of DISTAG disrupts basal cell fate, but also affects the differentiation medullary tissues. Thus, despite the deep evolutionary divergence and contrasting body plans of *Ectocarpus* and *Dictyota*, DISTAG retains a conserved role in basal development, while its analysis in *Dictyota* reveals an additional requirement for tissue differentiation and three-dimensional organisation. Our findings uncover a conserved developmental mechanism underlying distinct forms of brown algal multicellularity and establish a new approach for investigating development in morphologically complex brown algae.

## Introduction

Brown algae are one of the few eukaryotic lineages to have evolved complex multicellularity independently of animals, land plants and fungi, providing a powerful comparative system for investigating how developmental programmes arise and diversify (Coelho and Weijers, 2026; Denoeud et al., 2024). Brown algal body plans range from simple filaments to large, parenchymatous thalli with specialised tissues and developmental zones (Batista et al., 2024), yet the extent to which this morphological diversity is underpinned by conserved developmental mechanisms remains largely unknown. Most functional genetic studies have been performed in the filamentous brown alga *Ectocarpus*, leaving open whether developmental regulators identified in this experimentally tractable model have conserved functions in more morphologically complex brown algal lineages.

One such developmental process is the establishment of the apical-basal axis, an early and fundamental event in the development of complex multicellular organisms, including brown algae (Bogaert et al., 2022). In *Ectocarpus*, apical and basal cell identities are established during early development and ultimately give rise to distinct thallus and rhizoid systems. Forward genetic analysis identified DISTAG (DIS), which encodes the conserved TBCCd1 protein (André et al., 2013; Feldman and Marshall, 2009; Goncalves et al., 2010), as an important regulator of basal cell fate: *dis* mutants fail to establish a normal basal system and lose the characteristic bipolar pattern of germination (Godfroy et al., 2017). Whether DISTAG has a comparable developmental function in distantly related brown algae with more complex body plans is unknown.

The brown alga *Dictyota dichotoma* provides an attractive system in which to address this question. *Dictyota* belongs to the Dictyotophycidae, a lineage deeply divergent from the Fucophycidae, which includes *Ectocarpus*, and develops a more complex, parenchymatous thallus (Fritsch 1945; Gaillard et al.1986; Katsaros & Galatis, 1985). Its isomorphic diplohaplontic life cycle can be completed under laboratory conditions, and reproduction can be synchronised to obtain large numbers of gametes (Batista et al., 2024; Bogaert et al., 2016; Bogaert et al., 2023). Comparing DISTAG function between *Ectocarpus* and *Dictyota* therefore provides an opportunity to ask whether a developmental mechanism identified in a simple filamentous species has been conserved during the diversification of more complex brown algal body plans.

Such comparisons have been constrained by the limited availability of reverse genetic tools outside a small number of brown algal models. Genomic resources are now available across much of brown algal diversity (Denoeud et al., 2024; Liu et al., 2024; Lotharukpong et al., 2024; Vigneau et al., 2026) but functional approaches have not expanded at the same pace. More than half of the genes in surveyed algal genomes have no assigned function (Blaby-Haas and Merchant, 2019), leaving a substantial gap between gene discovery and functional understanding. This limitation is particularly apparent in developmental biology. Fucoid algae such as *Fucus* have long provided powerful cell biological systems for studying embryogenesis, cell polarity and axis formation (Brownlee et al., 2001; Coelho and Cock, 2020; Coelho et al., 2008) but their large size and long life cycles (Charrier et al., 2008; Yang, 2008) have made them poorly suited to genetic approaches. Conversely, reverse genetics has been developed primarily in smaller, laboratorytractable species such as *Ectocarpus*. Extending functional genetics to additional lineages and body plans is therefore essential for determining which developmental mechanisms are broadly conserved across brown algae and which have changed during the diversification of their multicellular forms (Coelho and Cock, 2020; Coelho et al., 2020).

CRISPR/Cas genome editing has begun to address this limitation, but in brown algae it has so far been restricted to members of the Fucophycidae, including *Ectocarpus*, *Scytosiphon* and the kelps *Saccharina*, *Laminaria* and *Undaria* (Badis et al., 2021; Martinho et al., 2026; Paix et al., 2026; Shen et al., 2023). Existing approaches exploit naturally cell wall-free stages, with preassembled Cas ribonucleoproteins (RNPs) introduced by microinjection or polyethylene glycol (PEG)-mediated delivery. Related DNA-free strategies have also enabled genome editing in other algae, including the green seaweed *Ulva* (Blomme et al., 2021; Ichihara et al., 2022; Ichihara et al., 2024). These approaches avoid the requirement for stable Cas expression but depend on reproductive or developmental stages that are not available or experimentally accessible in many brown algal lineages, limiting their broader application.

A second limitation is that the above mentioned genome-editing approaches target haploid cells. Haploid editing provides immediate access to recessive phenotypes but makes genes required for viability, growth or major developmental processes particularly difficult to investigate: strongly deleterious mutations may be lost before mutant individuals can be recovered, making it difficult to distinguish an essential gene from an unsuccessful editing event. Editing the diploid generation provides an alternative. Deleterious mutations can be maintained as heterozygous alleles and subsequently exposed in haploid progeny following meiosis, while segregation can separate the targeted allele from unlinked genetic changes. Establishing diploid editing would therefore broaden both the phylogenetic and functional scope of reverse genetics in brown algae.

Here, we use *Dictyota dichotoma* to test whether the developmental function of DISTAG has been conserved across deeply divergent brown algal lineages and contrasting multicellular body plans. We show that loss of DISTAG disrupts basal cell fate and rhizoid development in *Dictyota*, revealing a conserved requirement for DISTAG in basal development shared with *Ectocarpus*. In *Dictyota*, however, the consequences of DISTAG loss extend beyond early basal specification and persist during subsequent development of the parenchymatous thallus, indicating that this conserved regulator has acquired a broader developmental role in a more complex body plan. To enable this functional analysis, we exploit the reproductive biology of *Dictyota*, in which large, non-motile eggs are released synchronously and fertilised externally, creating a transient window between fertilisation and cell wall deposition (Bogaert et al., 2016; Bogaert et al., 2017b; Kumke, 1973). We use this window to establish a DNA-free CRISPR/Cas strategy in which preassembled Cas RNPs are delivered at fertilisation, directly targeting the newly formed diploid generation. Together, our results identify a conserved developmental mechanism underlying basal system development across distinct forms of brown algal multicellularity, while revealing how its developmental deployment has expanded in a more complex thallus, and establish fertilisation-based diploid editing as a new route for comparative developmental genetics in brown algae.

## Results

### DISTAG is conserved across deeply divergent brown algal lineages

*Dictyota* and *Ectocarpus* belong to two sister subclasses that separated deep in brown algal evolution, the Dictyotophycidae and Fucophycidae, respectively (Batista et al., 2024; Bringloe et al., 2020; Denoeud et al., 2024). *Dictyota* displays a more complex developmental organisation than the filamentous *Ectocarpus*. In both generations, the embryo of *Dictyota* polarises along an apical–basal axis (Bogaert et al., 2016; Bogaert et al., 2017a) (**Fig. 1A**). The basal cell gives rise to the rhizoid, whereas the apical cell generates an apical meristem cell which will produce a flattened parenchymatous thallus organised into morphologically distinct cortical and medullary cell layers (**Fig. 1A, B**). By contrast, the gametophyte and sporophyte generations of *Ectocarpus* are morphologically distinct and deploy different cell types and developmental programmes. The gametophyte develops an upright filament and rhizoids, whereas the sporophyte initially forms a prostrate filament, from which upright filaments and rhizoids subsequently arise (**Fig. 1A**). Therefore, *Dictyota* exhibits a more elaborate multicellular body plan, while *Ectocarpus* displays greater developmental differentiation between generations.

**Figure 1.**
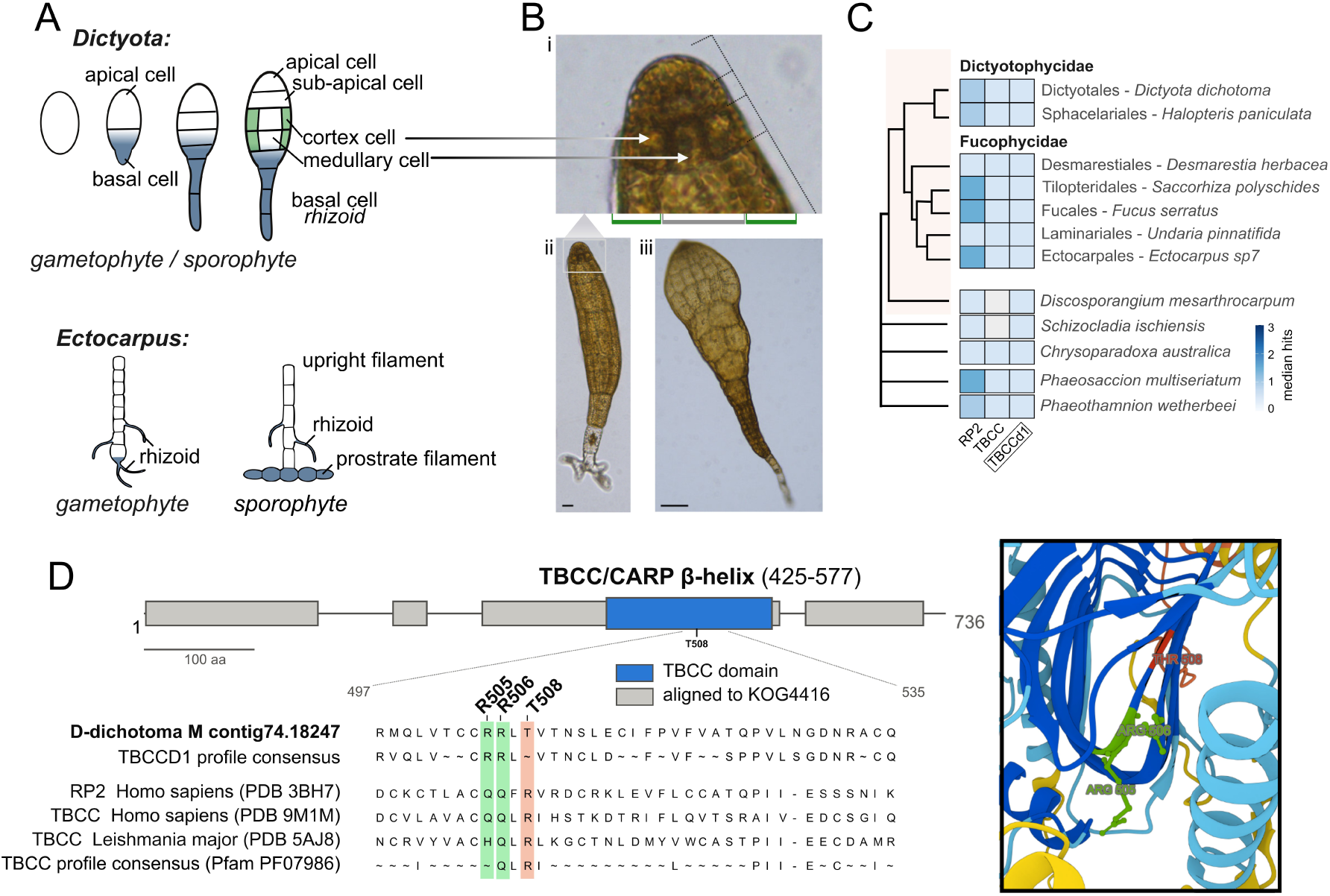
Divergent body plans but a conserved TBCCd1/DISTAG protein in *Dictyota* and *Ectocarpus*. **(A)** Schematic comparison of development in *Dictyota* (top) and *Ectocarpus* (bottom). In *Dictyota*, the first division of the zygote or spore establishes apical and basal cell lineages. The basal cell gives rise to the rhizoid, whereas the apical cell produces subapical cells that differentiate into cortical and medullary cell layers. The gametophyte and sporophyte follow the same developmental programme and are morphologically indistinguishable. In *Ectocarpus*, the gametophyte develops an upright filament and rhizoids, whereas the sporophyte initially forms a prostrate filament from which upright filaments and rhizoids subsequently arise. Blue, basal cell lineage and basal structures (rhizoid and prostrate filament); green, cortex; grey, medulla. **(B)** Patterning states of the embryo of *Dictyota*. (i) apical meristem of radially patterned embryo, dotted lines denote segmentation of the apical cell. Green sectors denote the (dividing) cortex layer, gray sector denotes the central (expanding) medullary cell. (ii) radially patterned embryo with basal rhizoid. Scale bar=20 µm (iii) young thallus following formation of the flattened blade. Scale bar=100 µm. **(C)** Copy number of the three TBCC-domain protein subfamilies across brown algae and related stramenopile classes. Rows represent taxonomic orders, with the number of genomes sampled indicated in parentheses; columns correspond to the RP2, TBCC and TBCCd1/DISTAG subfamilies. Cell colour indicates copy number in a representative genome from each order; copy numbers for all individual genomes are provided in Table S1. Brown algal orders are grouped by subclass (Dictyotophycidae and Fucophycidae), with Discosporangiales and non-Phaeophyceae classes shown separately. Proteins were identified by profile HMM searches of each proteome and assigned to the three subfamilies. **(D)** Domain organisation and structural features of *Dictyota* TBCCd1/DISTAG. Top, protein schematic showing regions aligned by HHpred to KOG4416 (grey) and the TBCC/CARP β-helix domain (blue). Consensus rows show the profile built from the 76 TBCCd1/DISTAG sequences identified here (59 from brown algae, 17 from other stramenopiles; Table S1). Bottom left, HHpred alignment of TBCCd1/DISTAG with RP2 and TBCC. R505 and R506 are indicated in green; a nearby arginine has been proposed to substitute for the missing arginine finger in TBCCd1 (Feldman and Marshall, 2009). Bottom right, AlphaFold 3 model of the β-helix domain coloured by pLDDT, with R505, R506 and T508 indicated.

The *Ectocarpus* genome encodes a single TBCCd1/DISTAG gene (Godfroy et al., 2017). To investigate its evolutionary conservation, we searched the predicted proteomes of 53 brown algal species spanning ten orders and both major subclasses, together with 14 additional stramenopiles representing ten classes. Searches were performed using profile hidden Markov models (HMMs) corresponding to the RP2, TBCC and TBCCd1/DISTAG subfamilies defined by Godfroy et al. (2017), and each identified sequence was assigned to its best-scoring subfamily (**Fig. 1C; Table S1**). TBCCd1/DISTAG homologues were widely distributed across the Phaeophyceae, including both Dictyotophycidae and Fucophycidae, as well as in most other stramenopile lineages examined. The gene was predominantly present as a single copy (**Fig. 1C; Table S1**), indicating broad evolutionary conservation with limited gene duplication.

The *Dictyota dichotoma* DISTAG protein also retained the characteristic structural features of the TBCCd1 family. Profile-based sequence comparisons identified the conserved TBCC/CARP β-helix domain, and structural modelling predicted the corresponding β-helical architecture (**Fig. 1D, Fig. S1**). In RP2 and TBCC proteins, this domain mediates GTPase-activating protein (GAP) activity through a conserved catalytic arginine finger (R118 in RP2 and R262 in TBCC; (Bartolini et al., 2002; Veltel et al., 2008)). Consistent with TBCCd1 homologues characterised in other eukaryotes (André et al., 2013; Feldman and Marshall, 2009), this canonical arginine is absent from *Dictyota* DISTAG and is replaced by a threonine (T508). However, two nearby arginine residues, R505 and R506, occupy positions corresponding to glutamine residues in RP2 and TBCC (**Fig. 1D**). Of these, R506 aligns with an arginine conserved in *Chlamydomonas* TBCCd1 (R418) and human TBCCd1 (R386), which has been proposed to functionally substitute for the missing catalytic arginine finger (Feldman and Marshall, 2009). The adjacent R505 residue is absent from these homologues, indicating a distinctive feature of the *Dictyota* protein. Thus, despite the deep evolutionary separation and pronounced differences in multicellular organisation between *Dictyota* and *Ectocarpus*, both species possess a conserved TBCCd1/DISTAG protein. We therefore asked whether this molecular conservation is accompanied by conservation of DISTAG function in basal development.

### Fertilisation-based CRISPR editing enables genome manipulation in diploid *Dictyota*

To determine whether the conserved *Dictyota* DISTAG protein retains its developmental function, we sought to generate *dis* mutants. However, existing PEG-mediated genome-editing approaches in brown algae rely on the manipulation of wall-less haploid reproductive cells (Martinho et al., 2026), and unfertilised *Dictyota* eggs do not normally develop into viable thalli. Moreover, editing in a haploid background immediately exposes recessive loss-of-function mutations, potentially limiting the recovery and propagation of mutants affecting essential developmental processes. We therefore developed an alternative strategy to introduce CRISPR reagents during fertilisation, allowing mutations to be generated directly in the diploid generation (**Fig. 2A,B**).

**Figure 2.**
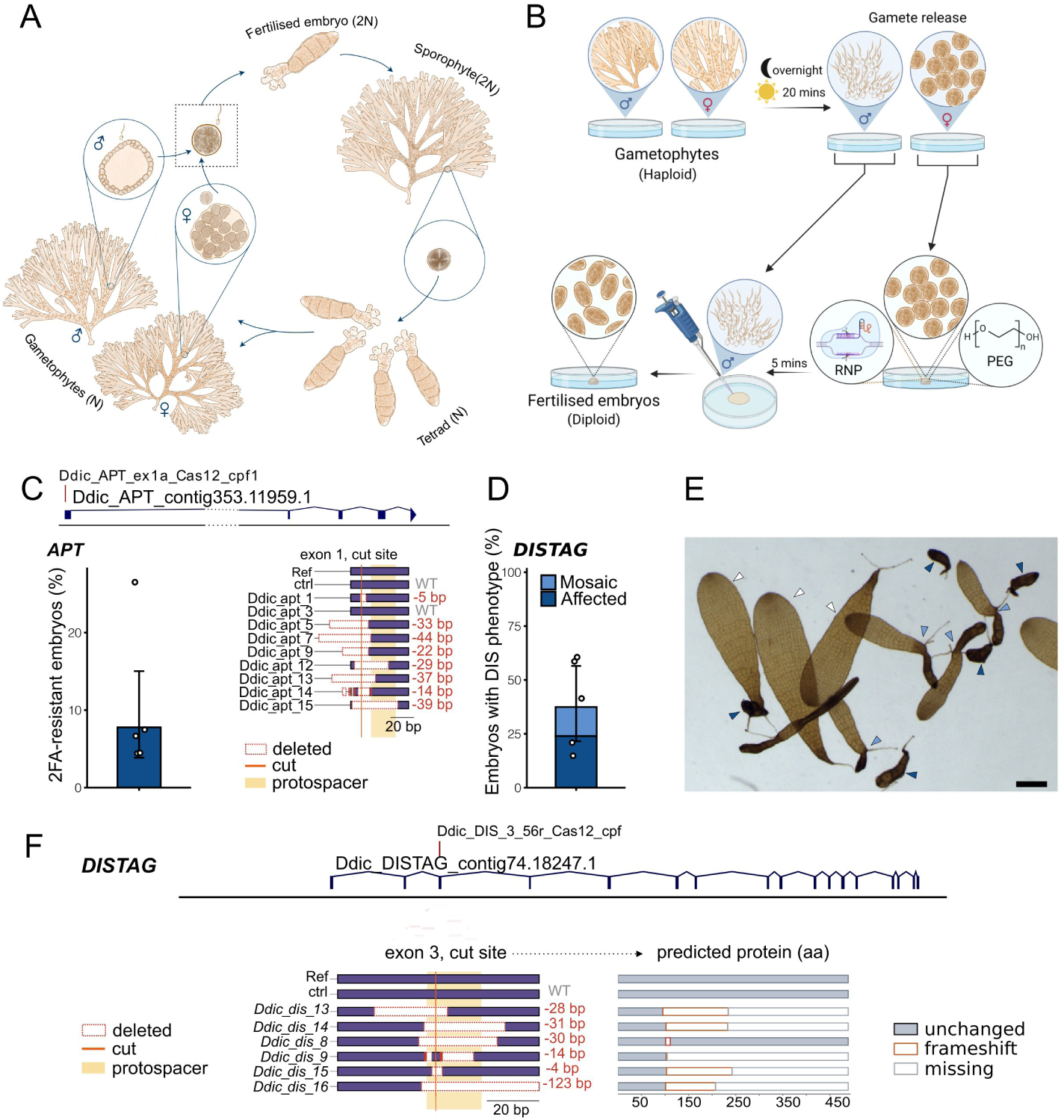
Fertilisation-based CRISPR–Cas12a editing generates diploid mutants in *Dictyota*. **(A)** Isomorphic diplohaplontic life cycle of Dictyota, showing haploid male and female gametophytes, gamete production, fertilisation, diploid sporophyte development and meiotic production of tetraspores. The stage targeted by RNP delivery is highlighted by a dotted box. **(B)** Experimental workflow for fertilisation-based genome editing. Eggs released from female gametophytes are incubated with preassembled Cas12a RNPs and PEG before sperm addition. Following fertilisation, treated zygotes are cultured and screened for edited phenotypes or selectablemarker resistance. **(C)** Validation of fertilisation-based editing by targeting APT. Frequency of 2-FA-resistant sporophytes following RNP treatment, together with representative mutations recovered at the APT target site. Genotyping confirmed APT mutations in 10/11 sequenced resistant individuals. The gene model is drawn to scale except for a 12,000-bp intronic region that was omitted for clarity and is indicated by a dotted line. **(D)** Frequency of abnormal sporophytes following DIS targeting (five plates, n = 445). Individuals were classified according to the extent of the developmental phenotype. Bars show model estimates with 95% confidence intervals; dots represent individual plates. **(E)** Representative wild-type (white arrow) and DIS-edited sporophytes illustrating the two principal mutant phenotypic classes: severely affected, dark, unflattened thalli (dark blue arrow) and individuals containing both abnormal and apparently normally developing regions (light blue arrow). Scale bar=500 µm. **(F)** Representative DIS alleles recovered from edited diploid sporophytes. The target sequence, predicted cleavage site, deletions and predicted protein consequences are indicated. Frameshift and in-frame mutations are distinguished.

*Dictyota* has an isomorphic diplohaplontic life cycle, in which haploid gametophytes and diploid sporophytes follow similar developmental programmes (**Fig. 2A**). Eggs are released without a cell wall and remain transiently accessible before fertilisation and the subsequent deposition of the zygotic cell wall (Bogaert et al., 2017). We reasoned that this brief window of accessibility could be exploited to deliver preassembled CRISPR–Cas12a ribonucleoprotein (RNP) complexes. To test this approach, freshly released eggs were incubated with RNPs and PEG for five minutes before the addition of sperm. Following fertilisation, the treated zygotes were diluted and cultured to obtain diploid sporophytes (**Fig. 2B; Methods**).

We first assessed the feasibility of this strategy by targeting *APT*, whose disruption confers resistance to 2-fluoroadenine (2-FA). We determined the sensitivity of *Dictyota* germlings to 2-FA and selected a concentration of 6.25 µM, which completely inhibited growth of untreated individuals (**Fig. S2**). Initial experiments in which selection was applied four to five days after fertilisation resulted in the death of all germlings (**Table S3**). We therefore delayed selection until two weeks after fertilisation, allowing the edited sporophytes to develop further before exposure to 2-FA. This delay may also have reduced the influence of maternally supplied APT products, although their persistence was not directly measured.

Under these conditions, 2-FA-resistant sporophytes were recovered in all five independent experiments, at frequencies ranging from 4.4% to 26.5% (pooled 6.7%, 83 resistant of 1,233 treated; binomial GLMM estimate 7.8%, 95% CI 2.9 to 16.7), whereas none of 254 untreated control individuals survived (**Fig. 2C; Table S3**). Sequencing of surviving sporophytes confirmed mutations at the *APT* locus in 10/11 individuals that were analysed (**Fig. 2C; Fig. S3**). These results demonstrate that CRISPR–Cas12a RNPs delivered during fertilisation can generate edited diploid sporophytes in *Dictyota*.

Having established the feasibility of fertilisation-based editing, we next targeted *DIS* to investigate the developmental function of the DISTAG protein. We initially co-delivered RNPs targeting *DIS* and *APT*, with the intention of enriching for edited individuals through 2-FA selection before screening for mutations at the *DIS* locus. Unexpectedly, conspicuous developmental abnormalities appeared in 15–61% of treated sporophytes before selection was applied (**Fig. 2D,E; Table S4**). The high frequency of these phenotypes suggested that *DIS* mutants could be recovered directly, without selectable-marker enrichment. We therefore screened treated individuals on the basis of their morphology and subsequently genotyped the *DIS* locus.

Two broad phenotypic classes were observed. The first comprised severely affected sporophytes that developed into small, dark thalli and failed to undergo the characteristic flattening observed in wild-type individuals (**Fig. 2E**). The second comprised sporophytes in which abnormal regions coexisted with apparently normally developing tissue, frequently including branches emerging from the basal region (**Fig. 2E**). We refer to the latter group as ‘mosaic’.

All severely affected individuals examined carried sequence variants at the *DIS* locus, five out of six of which were predicted to produce truncated DISTAG proteins (**Fig. 2F**). Sanger sequencing revealed no detectable wild-type sequence or evidence of heterozygosity in these individuals (**Fig. S4**), consistent with disruption of both alleles in at least a subset of the severely affected sporophytes. For the second class (referred to as ‘mosaic’), individuals exhibited overlapping sequencing traces downstream of the target site, indicating the presence of multiple sequence variants within the sampled tissue (**Fig. S4**). These patterns were consistent with heterozygosity or genetic mosaicism, although the two possibilities could not be distinguished from these data alone.

Together, these results establish fertilisation-based CRISPR editing as an effective strategy for generating diploid mutants in *Dictyota*. The recovery of individuals with distinct developmental phenotypes and allelic configurations also provided an opportunity to investigate whether disruptive *dis* alleles could be maintained and transmitted through the diploid generation.

### Diploid editing enables the recovery and transmission of recessive *DIS* mutations

The occurrence of sporophytes containing both abnormal and apparently normally developing tissues suggested that some edited individuals might retain functional *DIS* alleles in at least part of the thallus. We therefore investigated whether normally developing regions could be propagated to recover fertile individuals and assess the inheritance of edited *dis* alleles.

During prolonged cultivation of 36 initially severely affected sporophytes, 25 produced one or more branches with apparently wild-type morphology. These branches were not restricted to the basal region, indicating that normal development could emerge from different parts of an abnormal thallus. We excised and propagated the normally developing branches, removing the deteriorating abnormal tissue (**Fig. 3A**). The resulting subcultures developed largely normally, and 20 of the 25 subsequently became fertile, allowing us to investigate the transmission of edited *DIS* alleles through meiosis.

**Figure 3.**
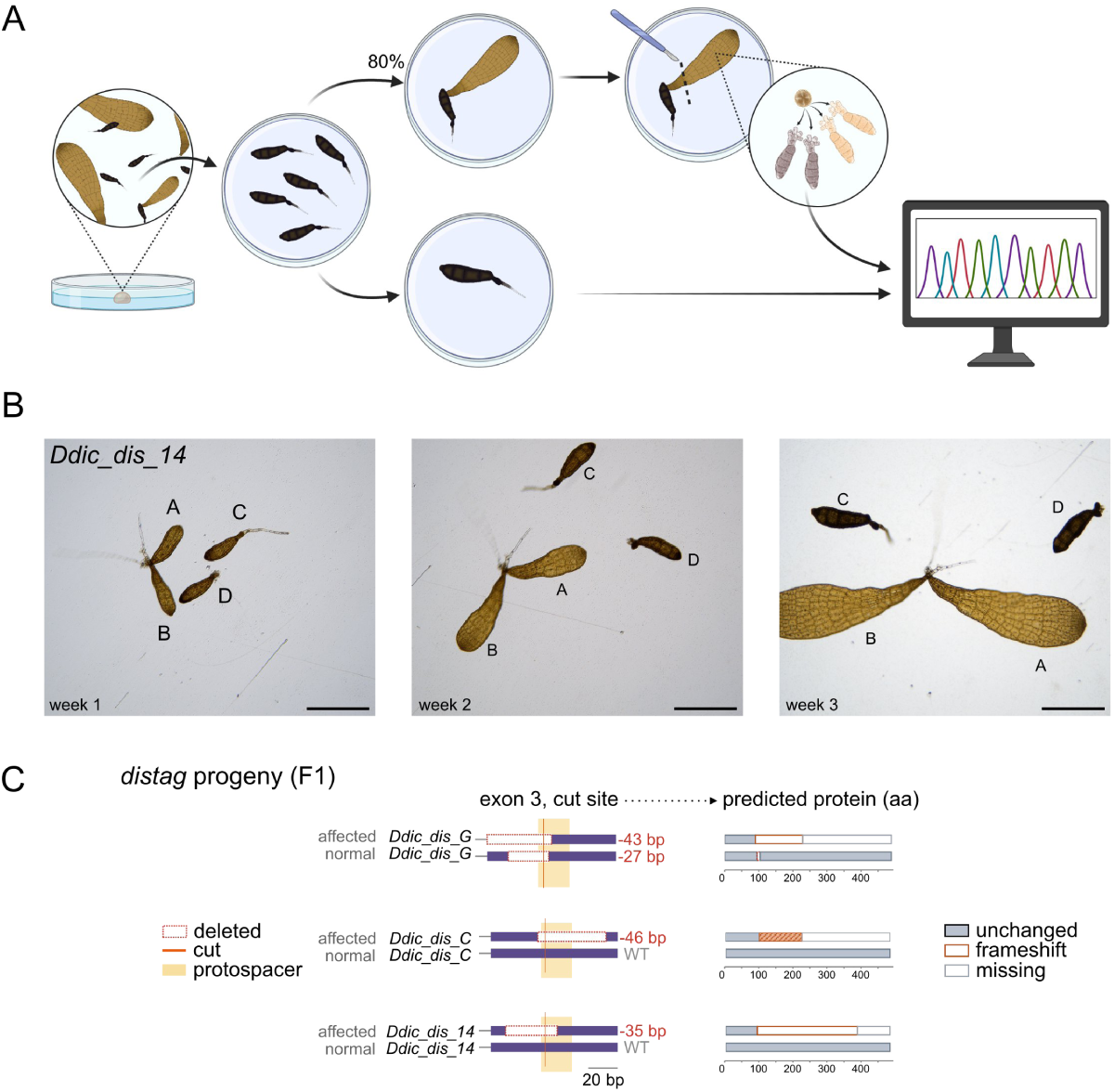
Recovery and meiotic transmission of recessive *dis* mutations. **(A)** Experimental strategy for recovering heritable dis alleles. Edited sporophytes exhibiting severe developmental abnormalities were maintained in culture. Normally developing branches were isolated and propagated to obtain fertile sporophytes, from which haploid progeny were recovered for phenotypic and genetic analysis. Of 36 individuals maintained in culture, 25 produced normal branches, 20 of these became fertile, and 15 produced both normal and abnormal haploid progeny..**) (B)** Developmental phenotypes of haploid progeny recovered from edited diploid parents. Representative normal and mutant siblings are shown during development, illustrating the progressive appearance of the dis phenotype. In the photos, A-D indicate always the same followed individuals Scale bars=500 µm. **(C)** Association between DIS genotype and developmental phenotype in haploid progeny from two edited parental sporophytes. Target-site sequences, deletions and predicted protein consequences are shown for normal and abnormal progeny. Disruptive mutations are associated with abnormal development.

Analysis of their haploid progeny revealed two clearly distinguishable phenotypic classes. Most fertile parental sporophytes (15/20) produced both normally developing progeny and progeny displaying severe developmental abnormalities resembling those observed in strongly affected diploid individuals (**Fig. 3B,C, Table S5**). Although mutant and normal progeny were initially similar, developmental differences became progressively apparent over the following weeks. Mutant individuals became darker, failed to flatten and showed strongly impaired thallus development, whereas their normal siblings continued to develop the characteristic blade morphology (**Fig. 3B**).

Genotyping revealed a consistent association between developmental phenotype and *DIS* genotype (**Fig. 3C**). Abnormal progeny carried disruptive mutations in *DIS*, whereas normally developing progeny carried either the wild-type allele or sequence variants predicted to preserve protein function. Where complete tetrads could be recovered, the two phenotypic classes segregated in a 1:1 ratio (**Fig. 3B**), consistent with Mendelian segregation at a single nuclear locus. Together with the genotype–phenotype association and the recovery of similar abnormalities from independently edited individuals, these results provide strong genetic evidence that disruption of *DIS* is responsible for the developmental defects.

The recovery of both mutant and normally developing haploid progeny from individual diploid parents further demonstrated that edited *dis* alleles introduced during fertilisation could be transmitted through meiosis. These segregation patterns were consistent with the presence of functional and disrupted *DIS* alleles in the reproductive lineage of the parental ‘mosaic’ sporophytes. The results demonstrate that disruptive alleles can be maintained in diploid individuals retaining functional *DIS* copies.

Haploid *dis* gametophytes displayed developmental abnormalities similar to those observed in affected diploid sporophytes, whereas wild-type individuals developed similarly in both generations (**Figs. 2E, 3B**). Thus, loss of *DIS* affects development in both generations of the isomorphic *Dictyota* life cycle.

Although haploid *dis* gametophytes formed reproductive structures, attempted crosses between male and female mutant individuals did not yield detectable zygotes. This suggests that loss of *DIS* may also interfere, directly or indirectly, with successful sexual reproduction. Consequently, homozygous zygotes could not be established and studied through these crosses, and disruptive *dis* alleles were instead maintained in diploid *DIS/dis* individuals retaining functional alleles.

The segregation of normal and mutant phenotypes supports the recessive nature of the *dis* mutations: a functional *DIS* allele is sufficient to support essentially normal development, whereas haploid progeny carrying disruptive alleles express the mutant phenotype. Thus, fertilisation-based diploid editing provides a means of maintaining deleterious developmental mutations and recovering their phenotypic consequences following meiosis (**Fig. 3**).

### DISTAG is required for basal cell identity and medullary differentiation in *Dictyota*

To investigate the developmental function of DISTAG in *Dictyota*, we compared wild-type and *dis* mutants throughout development, focusing on the establishment of basal structures and the organisation of the multicellular thallus. In wild-type *Dictyota*, development begins with an asymmetric division of the zygote or spore that establishes the apical–basal axis (Bogaert et al., 2017a). The basal cell gives rise to the primary rhizoid, whereas the apical region establishes the growing thallus. Subsequent divisions generate a radially organised young thallus, which progressively differentiates into a flattened blade composed of an outer cortex of small, chloroplast-rich cells surrounding an inner medulla of larger, chloroplast-poor cells (**Fig. 1A**).

During the earliest stages of development, *dis* mutants were morphologically similar to wild-type individuals (**Fig. 3B**), paralleling the late emergence of the *dis* phenotype in edited sporophytes, and which could be explained by the presence of deposited DIS protein in the spore or egg. Interestingly, DISTAG transcripts were low in gametes and increased progressively during early development (**Fig. S2A**). However, clear developmental abnormalities became apparent as growth proceeded. In wild-type individuals, the primary rhizoid elongated and maintained its characteristic morphology, and additional rhizoids developed from the basal region. In contrast, although *dis* mutants initially produced a primary rhizoid, this structure subsequently became swollen and pigmented, progressively developing into a chain of parenchymatous, chloroplast-containing cells (**Fig. 4A**). Moreover, no additional rhizoids were observed. These results indicate that DISTAG is required for the maintenance of rhizoid identity and normal development of the basal attachment system, consistent with its previously described function in *Ectocarpus*.

**Figure 4.**
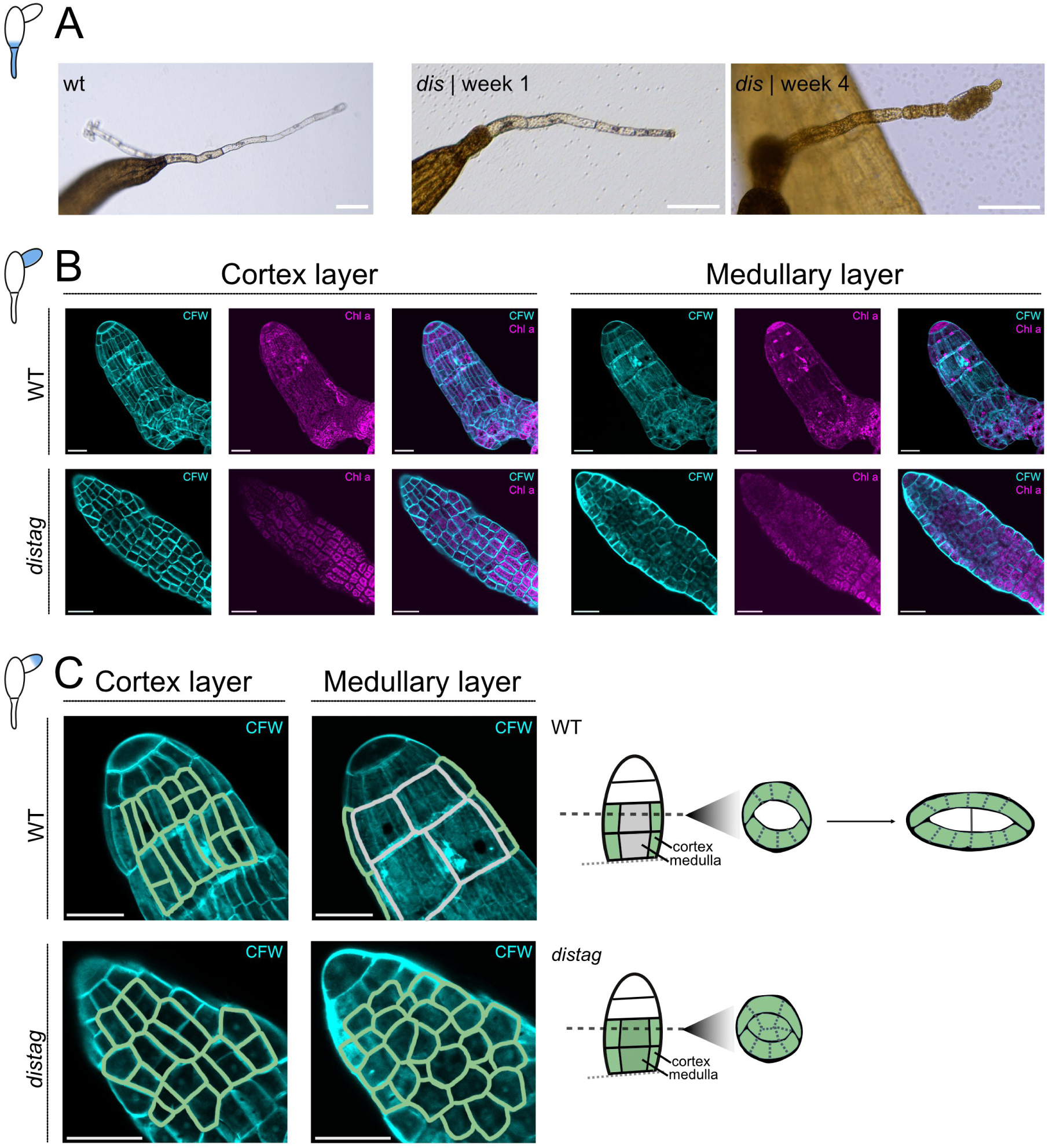
DISTAG is required for rhizoid identity and medullary tissue differentiation in *Dictyota*. (**A)** Development of the primary rhizoid in wild-type and *dis* individuals at one and four weeks. In wild type, the primary rhizoid maintains its elongated morphology and additional rhizoids develop. In *dis* mutants, the primary rhizoid initially elongates but subsequently becomes swollen and pigmented, forming a chain of parenchymatous cells. No additional rhizoids are produced. Scale bars=100 µm. **(B)** Confocal optical sections through the cortical (left) and internal medullary (right) regions of wild-type (top) and *dis* (bottom) thalli. Cell walls are stained with Calcofluor White (cyan), and chloroplast autofluorescence is shown in magenta. In wild type, the internal region contains large, chloroplast-poor medullary cells, whereas the corresponding region of *dis* thalli contains numerous smaller, chloroplast-rich cells. Scale bars=50 µm. **(C)** Left, cell outlines traced from the optical sections shown in (B), highlighting the differences in cortical and internal tissue organisation between wild-type and *dis* thalli. Cortical cells are shown in green and medullary cells in grey. Right, schematic representation of thallus organisation and development, adapted from Katsaros and Galatis (1985). In wild type, the central medulla differentiates into enlarged cells and the thallus develops into a flattened blade. In *dis* mutants, internal cells retain a smaller, cortex-like morphology, and the thallus fails to flatten. Rhizoids are shown in blue. Scale bars, 50 µm.

The developmental defects of *dis* mutants were not restricted to the basal region. Whereas wildtype thalli progressively flattened to form the characteristic blade, mutant thalli remained narrow, darkly pigmented and approximately radially symmetric, with strongly altered internal tissue organisation (**Fig. 4B,C**). To investigate the cellular basis of this phenotype, we examined cortical and internal tissues using confocal microscopy and Calcofluor White staining. These analyses were performed on later-developing branches, allowing us to characterise the organisation of newly formed tissues beyond the earliest stages of development.

In wild-type thalli, the cortex consisted of small, chloroplast-rich cells surrounding a central medulla composed of larger, chloroplast-poor cells. In contrast, the central region of *dis* thalli contained numerous smaller, chloroplast-rich cells, which resembled cortical cells rather than differentiated medullary cells (**Fig. 4B,C**). Although these internal cells were not identical in size to typical cortical cells, the characteristic enlarged medullary cells of wild-type thalli were absent. Thus, loss of DISTAG disrupts the normal differentiation of internal tissues, suggesting that cells occupying the medullary region fail to acquire or maintain a fully differentiated medullary identity.

The altered organisation of internal tissues was accompanied by a failure to establish the flattened blade morphology characteristic of wild-type *Dictyota*. These observations suggest that normal medullary differentiation and blade morphogenesis are closely associated, although the present data do not establish whether the failure of medullary differentiation directly causes the morphological defect or whether both arise from an earlier disruption of thallus development.

Similar developmental abnormalities were observed in haploid *dis* gametophytes and diploid *dis* sporophytes, consistent with the largely isomorphic development of the two generations. DISTAG is therefore required for normal thallus development in both phases of the *Dictyota* life cycle.

Together, these results reveal both conserved and broader developmental functions of DISTAG in brown algae. As previously described in *Ectocarpus*, DISTAG is required for normal basal development in *Dictyota*, suggesting conservation of this function across divergent brown algal lineages. However, in *Dictyota*, loss of DISTAG also disrupts internal tissue differentiation and the formation of the flattened blade. These findings implicate DISTAG in the coordination of cell identity and tissue organisation during the development of a complex multicellular thallus.

## Discussion

Our study shows that DISTAG, a developmental regulator first characterised in the filamentous brown alga *Ectocarpus*, retains a conserved role in basal development in the distantly related and morphologically distinct *Dictyota*. Despite their contrasting body plans, loss of DISTAG affects rhizoid development in both species, suggesting that this function predates the divergence of the Dictyotophycidae and Fucophycidae. In *Dictyota*, *dis* mutants initially produce a primary rhizoid, which subsequently becomes swollen, pigmented and parenchymatous, while additional rhizoids fail to develop. Similar defects occur in haploid gametophytes and diploid sporophytes, as previously observed in *Ectocarpus* (Godfroy et al., 2017). These findings indicate that the role of DISTAG in basal development has been maintained across divergent brown algal body plans and both generations of the life cycle.

In *Dictyota*, DISTAG disruption also affects aspects of development that have not been associated with this regulator in *Ectocarpus*. Mutant thalli fail to acquire the characteristic flattened blade morphology and instead retain an approximately radial symmetric organisation. They also lack the enlarged, chloroplast-poor medullary cells characteristic of wild-type thalli, containing instead numerous small, chloroplast-rich cells. Whether these abnormalities result from altered cell division patterns, defects in cell differentiation or a combination of both remains unclear. Nevertheless, these observations extend the developmental functions associated with DISTAG beyond basal development and suggest that its contribution to tissue organisation may differ between brown algal body plans.

The molecular mechanisms underlying DISTAG function remain unclear. DISTAG belongs to the TBCCd1 family and retains the characteristic TBCC/CARP β-helix fold, but lacks the canonical catalytic arginine required for the GTPase-activating activity of TBCC and RP2 (Bartolini et al., 2002; Veltel et al., 2008). Instead, *Dictyota* DISTAG conserves an alternative arginine (R506), proposed to substitute for the missing catalytic residue in *Chlamydomonas* and human TBCCd1 proteins, together with a neighbouring arginine (R505) of unknown significance (Feldman and Marshall, 2009). TBCCd1 proteins have been implicated in centrosome and cytoskeletal organisation in other eukaryotes (Feldman and Marshall, 2009; Gonçalves et al., 2010; André et al., 2013), and *Ectocarpus dis* mutants exhibit defects in nuclear and centriole positioning, Golgi organisation and microtubule architecture (Godfroy et al., 2017). These observations suggest that the developmental role of DISTAG may be linked to a more broadly conserved cellular function.

The progressive appearance of the *dis* phenotype also raises questions about the timing of DISTAG function. Mutants initially establish apical-basal polarity and produce a primary rhizoid before developing pronounced abnormalities. This delay could reflect the persistence of maternally supplied DISTAG products or a requirement for DISTAG during later developmental stages. Distinguishing between these possibilities will require analysis of DISTAG protein localisation and dynamics during early development, for which tools such as endogenous fluorescent tagging are not yet available in *Dictyota*.

A second important outcome of this study is the establishment of genome editing directly in the diploid generation of *Dictyota*. Previous DNA-free genome-editing approaches in macroalgae have predominantly targeted haploid reproductive stages, including gametes and meiospores (Ichihara et al., 2022; Blomme et al., 2025; Martinho et al., 2026; Paix et al., 2026). Although these approaches allow mutant phenotypes to be characterised directly, recessive deleterious mutations are immediately exposed and may prevent the recovery or propagation of mutant individuals. Editing the diploid generation offers a complementary strategy, allowing disruptive alleles to be maintained in heterozygous individuals and subsequently recovered in haploid progeny following meiosis.

The inheritance of *DIS* mutations illustrates this advantage. Edited sporophytes retaining functional *DIS* alleles developed sufficiently normally to reach reproductive maturity and transmit disruptive alleles to their haploid progeny. Normal and mutant phenotypes co-segregated with functional and disruptive *DIS* alleles, respectively, and complete tetrads showed 1:1 segregation, consistent with Mendelian inheritance at a single nuclear locus (**Fig. 3C, Table S5**). In contrast, severely affected diploid individuals with evidence of disruption of both *DIS* alleles failed to produce progeny. These results support the recessive nature of the *dis* phenotype and demonstrate how diploid editing can facilitate the recovery and maintenance of mutations affecting development.

The reproductive biology of *Dictyota* is particularly well suited to this approach. Eggs are released without a cell wall and remain accessible during a brief period around fertilisation, before deposition of the zygotic wall. Delivery of preassembled CRISPR–Cas ribonucleoproteins during this window allowed the generation of edited diploid sporophytes without stable expression of genome-editing components. To our knowledge, this is the first application of fertilisation-based genome editing in algae and the first direct CRISPR editing of the diploid sporophyte generation in a brown alga.

One limitation of editing during fertilisation is the generation of heterozygous and mosaic individuals, which can complicate genotype–phenotype associations. This can be resolved within the edited generation by allowing some additional weeks for vegetative growth and isolating clonally uniform branches before genotyping. In our study, propagation of normally developing branches, followed by genotyping and analysis of meiotic segregation, allowed the inheritance of edited alleles to be resolved. Mosaicism may also allow multiple independently edited alleles to be recovered from individual sporophytes, while heterozygosity provides a means of maintaining recessive deleterious mutations.

The efficiency of fertilisation-based editing makes this approach particularly useful for functional studies. Following *APT* targeting, a range of 4-26.5% of treated sporophytes became resistant to 2-FA, whereas 15–61% of sporophytes targeted at *DIS* displayed mutant phenotypes. These frequencies compare favourably with the 4.5–8.6% 2-FA resistance reported following microinjection of *Saccharina* gametophytes (Shen et al., 2023) and with reported outcomes of PEG-mediated editing in other brown algae (Martinho et al., 2026), although differences in experimental conditions and scoring methods preclude direct comparisons. Importantly, fertilisation-based editing can be applied to batches of zygotes rather than individual cells, and the frequency of *DIS* phenotypes allowed mutants to be identified by direct phenotypic screening and genotyping, without selectable-marker enrichment.

Although the large, externally fertilised eggs of *Dictyota* provide particularly favourable conditions, the underlying principle may be applicable to other algae. Brown algae exhibit considerable diversity in their reproductive biology, and different species will likely require different strategies for introducing genome-editing reagents. Exploiting transient periods of cellular accessibility during reproduction or early development may therefore provide an alternative route to functional genetics in species that remain difficult to manipulate.

Together, our results show that DISTAG has retained a role in basal development across deeply divergent brown algal lineages, despite substantial differences in their multicellular organisation. The defects in basal cell differentiation and in medullary cell differentiation may be explained by the proposed function of TBCCd1 in positioning the nucleus, Golgi and microtubule-organising centre, reflected in the altered Golgi organisation, disrupted microtubule networks and aberrant nuclear positioning in *Ectocarpus* dis mutants (Godfroy et al., 2017) and as centriole positioning defects in *Chlamydomonas* TBCCd1 mutants (Feldman and Marshall, 2009). The additional defects in tissue organisation raise the possibility that conserved developmental regulators contribute to different aspects of morphogenesis in different body plans. Whether these differences reflect evolutionary changes in DISTAG function or distinct developmental consequences of disrupting a conserved cellular process remains unresolved. Comparative functional studies across brown algae will help determine how conserved developmental mechanisms have contributed to the diversification of their multicellular body plans.

## Material and Methods

### Algal material and culture

Experiments were conducted using unialgal laboratory cultures of *Dictyota dichotoma* (Hudson) J.V.Lamouroux. The strains used were KB07f (female) and KB07m (male), collected near the Roscoff Biological Station in Brittany, France, and ODC_1387m (male), collected at l’Ancien Fort Croy in Wimereux, France. Gametophytes were cultured in aerated flasks or crystallising dishes at approximately 20 °C, under 10 µmol m^-2^ s^-1^ light intensity with a 16:8 light:dark photoperiod. Fertility and growth were induced under the same temperature and light conditions, with an overnight light exposure once per month to simulate natural moonlight and enhance reproductive synchronisation. Under this regime gametogenesis followed a semilunar rhythm, with fertile thalli obtained approximately 10 to 14 days before and after full moon. All cultures were grown in filtered natural seawater (NSW), sterilised by autoclaving, and enriched with half-strength Provasoli enriched seawater (PES). Media in bubbling flasks was refreshed weekly to support optimal algal growth.

### 2-FA pharmacology assay

The 2-FA sensitivity of wild-type gametophytes was tested for *D. dichotoma*. Gametophytes were grown in 24-well plates for 2 weeks before 2-FA was added. The gametophytes were exposed to different concentrations of 2-FA mixed with equal volumes of NSW+PES (n=3) at the following concentrations (μM): (0.05, 0.1, 0.2, 0.39, 0.78, 1.56, 3.13, 6.25, 12.5). Controls of NSW and DMSO were used. The minimum lethal dose was determined to be 6.25 μM after 2 weeks by the concentration at which 25 randomly selected blades of D. dichotoma per well showed signs of cell death. This concentration of 2-FA was used for future gene editing procedures.

### Sequence comparison

Genome assemblies and annotations of stramenopile (53 of them Phaeophyceae) strains were taken from Denoeud et al. 2024. Missing protein sets were regenerated from the GFF and assembly with gffread (Pertea and Pertea, 2020).

A profile HMM built with hmmbuild from the TBCC-domain alignment of Godfroy et al. (2017; 25 sequences of the RP2, TBCC and TBCCd1/DISTAG subfamilies) was searched against each proteome with hmmsearch (HMMER; Eddy, 2011; full-sequence E-value <= 1e-5). Hits were assigned to a subfamily by hmmscan against three subfamily profiles built from the same alignment, using the best-scoring profile; assignments in which the two best profiles differed by less than 5 bits were flagged as ambiguous (**Table S1**).

Domain organisation was analysed with HHpred (Steinegger et al., 2019) in the MPI Bioinformatics Toolkit (Zimmermann et al., 2018), using an alignment of the 76 TBCCd1/DISTAG sequences with the *D. dichotoma* protein as query, against PDB_mmCIF70, Pfam-A v38.2, NCBI Conserved Domains v3.19, SMART v6.0, COG/KOG v1.0, ECOD F70 and CATH S40 v4.3 with default settings. The full-length protein was modelled with AlphaFold 3 (Abramson et al., 2024). The betahelix (residues 425-577) was compared with *Chlamydomonas reinhardtii* (UniProt C6EWZ9) and human TBCCd1 (Q9NVR7) by local pairwise alignment (BLOSUM62).

### RNP assembly

The genes of APT and DISTAG were identified in *Dictyota dichotoma* by searching the *Ectocarpus* sp. 7 protein sequences against the *D. dichotoma* predicted proteome with blastp (BLAST+ v2.13.0; Camacho et al., 2009). A single gene was found for both APT on chromosome 28 (mRNA_D-dichotoma_M_contig353.11959.1) and DISTAG on chromosome 30 (mRNA_D-dichotoma_M_contig74.18247.1) Guides were created using CRISPOR (Concordet and Haeussler, 2018) (**Table S2**). LbCas12a crRNAs (100 μM, IDT, Leuven, Belgium) were complexed individually by combining 4 μl crRNA (30 μM, IDT), 8 μl LbCas12a protein (15.6 μM, IDT), and 8 μl of 2.5X NEB 3.1 Buffer (New England Biolabs, Ipswich, MA, USA) diluted in IDTE Buffer (pH 7.5, IDT) to a final volume of 20 μl, and incubating at room temperature for 15 minutes. This RNP complex mixture was prepared the day before the experiment for each guide and was stored at 4°C until use.

### Transfection and Fertilisation

Fertile male and female gametophytes were sorted separately and placed in plastic Petri dishes (90 x 15mm) and were kept in darkness overnight in a sealed container to simulate nocturnal conditions. The following morning, gametophytes were exposed to light and transferred into fresh medium. After approximately 20 minutes of light exposure at room temperature, female gamete release occurs and the released gametes are collected for subsequent experiments. Male gametes are released in the same way from gametophytes for subsequent experiments. The transfection was mediated with PEG treatment where 100 μl of female gametes were pipetted into a plastic petri dish (90×20mm) along with 5μl of RNP complex for each guide (or a control solution using IDT buffer) and pre-treated with 40% w/v PEG-8000 (Fisher Scientific, Schwerte, Germany) in a 1:1 volume ratio with the combination of female gametes and RNP. This pretreatment was incubated at room temperature for 5 minutes before freshly released male gametes were added to induce fertilisation. This mixture of RNP, PEG, and gametes was further incubated at room temperature for 10 min to maximise the transient accessibility of the eggs. Following this incubation, 20ml of half strength PES medium was added to the petri dish. The dishes were then placed in darkness overnight and then returned to normal culture conditions. Two weeks after transfection, when the embryos had flattened from the initial radial stage, 2-FA was added. Resistant individuals were isolated after 10 days of selection.

### Genotyping

Genotyping PCR was performed using the Terra PCR Direct Polymerase Mix (Takara) using 2 μL of a suspension of small fragments of algal tissue (approximately 1 mm^2^) ground in 20 μL of Nuclease free Water (Qiagen). Sanger sequencing was performed by Azenta. Target regions of APT and DIS were amplified with the primers listed in **Table S2**.

Sanger traces of sequenced lines and the untreated control were analysed with Biopython v1.84, NumPy v2.4.6 and Matplotlib 3.11.2. Reads were quality trimmed with the modified Mott algorithm using Phred quality values using error probability limit (error limit 0.01, approx. Q20; Ewing and Green, 1998). Reads were aligned to the reference amplicon by local Smith-Waterman alignment using Biopython. Called alleles were translated with Biopython (standard genetic code).

### Editing efficiency

Each Petri dish was treated as one independent transformation. Editing efficiency at APT was calculated as the number of embryos surviving 2-FA selection divided by the number of embryos treated, and at DIS as the number of embryos displaying a mosaic or fully mutant phenotype divided by the number of embryos treated. Proportions were modelled with a binomial generalised linear mixed model with a logit link and dish as a random intercept, fitted with glmer in lme4 in R; bars show the back-transformed fixed-effect intercept, error bars the 95% Wald confidence interval and points the observed proportion per dish, with the DIS bar subdivided into mosaic and fully mutant fractions.

### Expression analysis

Expression of APT and DISTAG was extracted from the *D. dichotoma* developmental RNA-seq dataset of (Bogaert et al., 2026), comprising female gametes, zygotes, and polarisation, asymmetric cell division, four-celled and patterning stages. Counts were normalised with DESeq2 and stages compared with female gametes by Wald test with Benjamini-Hochberg correction.

### Confocal Microscopy

Cell walls were stained with 0.001% (w/v) Calcofluor White in filtered natural seawater and imaged directly using a Leica SP8 upright confocal microscope equipped with a 40× water-immersion objective. Calcofluor White was excited at 405 nm, and fluorescence emission was collected between 420 and 480 nm. Chloroplast autofluorescence was recorded in a separate channel using excitation at 638 nm and emission detection between 660 and 690 nm. Images were acquired as Z-stacks with a step size of 0.48 µm. Laser power was adjusted using Z-compensation to account for signal attenuation with imaging depth, applying the same settings to wild-type and mutant samples. All other acquisition parameters were kept constant, and images were checked for signal saturation. Single optical sections are presented in the figures.

### Image Analysis

Images were analysed using Fiji/ImageJ version 1.54p. Image adjustments were restricted to linear brightness and contrast adjustments, applied uniformly across each entire panel. Images presented in the figures are confocal optical sections. For Fig. 4C, cell outlines were manually traced using the Calcofluor White channel. Cells were classified as cortical or medullary based on their position within the tissue, differences in cell size and chloroplast autofluorescence.

## Acknowledgments

We thank Daniela (Lulo) Gallego Calderon for the algal drawings and illustrations. We thank Andrea Belkacemi for assistance with algal cultures. This study was funded by the Max Planck Society, the European Research Council grant 864038 (SMC), the Bettencourt Foundation (SMC) and the Moore Foundation (SMC).

## Data availability

All strains used in this study can be distributed upon contacting the corresponding authors.

## Contributions

KB and SMC conceived and designed the experiments; KB supervised the project. AK, MR and KB performed experiments. AK and KB curated the data. AK and KB wrote the original draft with support from SMC. SMC reviewed and edited the manuscript. All authors read and approved the final manuscript.

## Supplemental figures

**Figure S1.**
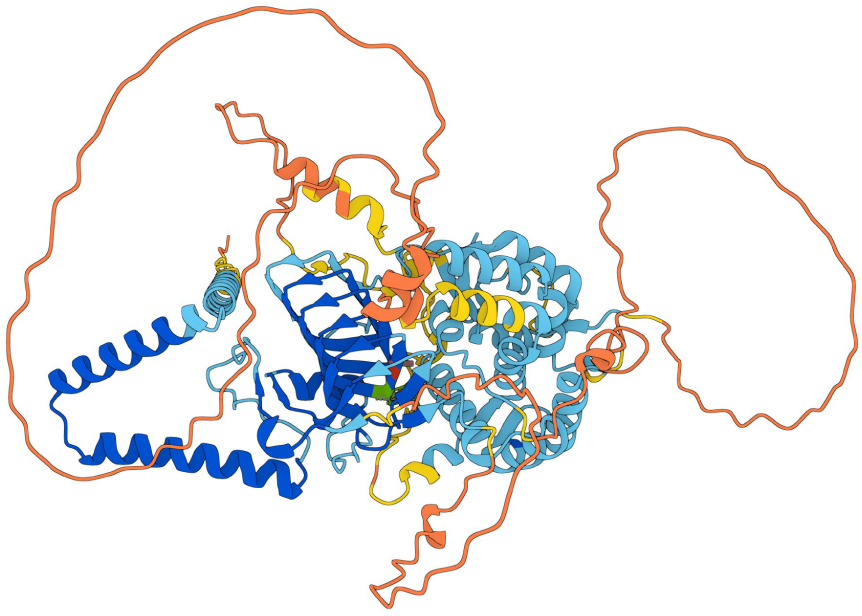
AlphaFold 3 model of full-length *Dictyota* TBCCd1/DISTAG, coloured by per-residue confidence (pLDDT): dark blue, very high (>90); light blue, confident (70-90); yellow, low (50-70); orange, very low (<50). The TBCC/CARP β-helix domain (residues 425-577) is predicted with high confidence and packs against an α-helical domain. T508 (red), which sits where the canonical arginine finger would be, and R505 and R506 (green) are shown on the v-helix.

**Figure S2.**
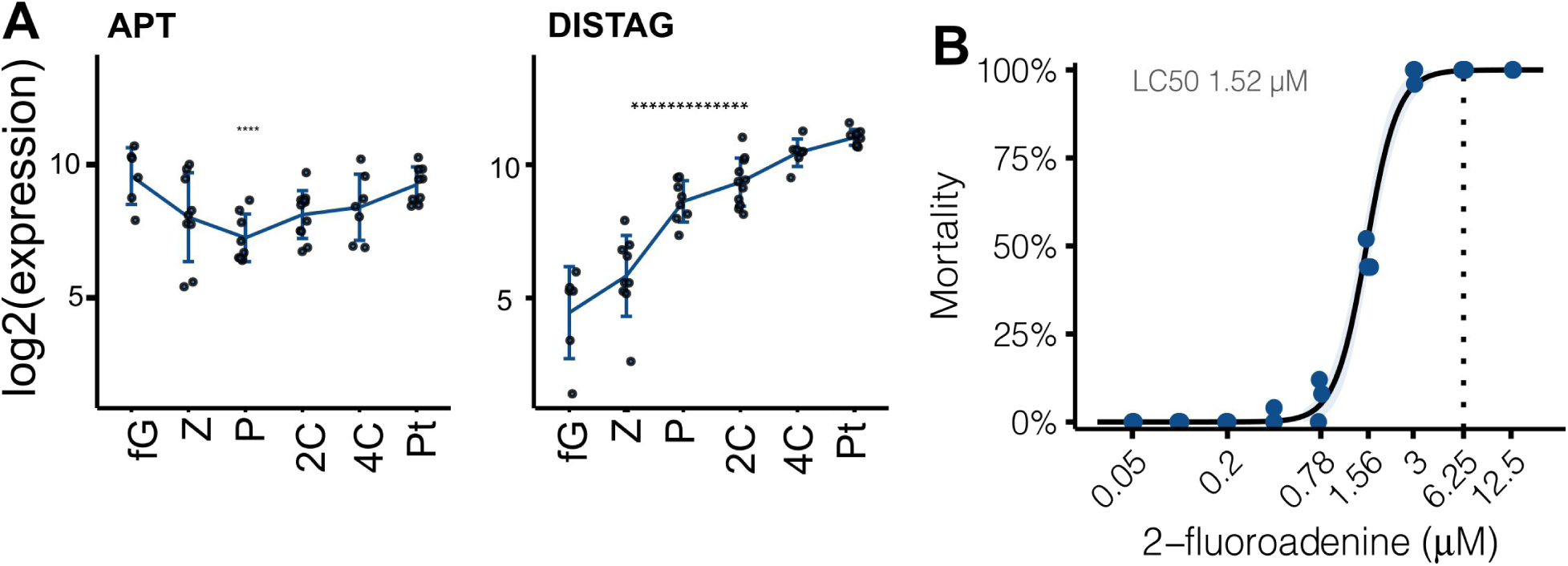
APT and DISTAG expression during early development, and 2-fluoroadenine sensitivity of *Dictyota* germlings. **(A)** Expression of APT (D-dichotoma_M_contig353.11959) and DISTAG (D-dichotoma_M_contig74.18247) across early development: female gametes (fG), zygote (Z), polarisation (P), asymmetric cell division (2C), four-celled (4C) and patterning (Pt). Points, individual replicates; line, mean per stage; error bars, ± SD; y axis, log2(DESeq2-normalised counts + 1). Asterisks mark stages differing from female gametes (DESeq2 Wald test, Benjamini-Hochberg adjusted p < 0.05 and |log2FC| ≥ 1; *p < 0.05, **p < 0.01, ***p < 0.001). APT is already abundant in gametes and the early zygote, whereas DISTAG is low in gametes and rises during development. **(B)** Mortality of wildtype germlings after 2 weeks in 2-fluoroadenine. Points, observed mortality; curve, fitted dose-response model with 95% confidence band; LC50 = 1.52 µM. Dotted line, the 6.25 µM used for selection.

**Figure S3.**
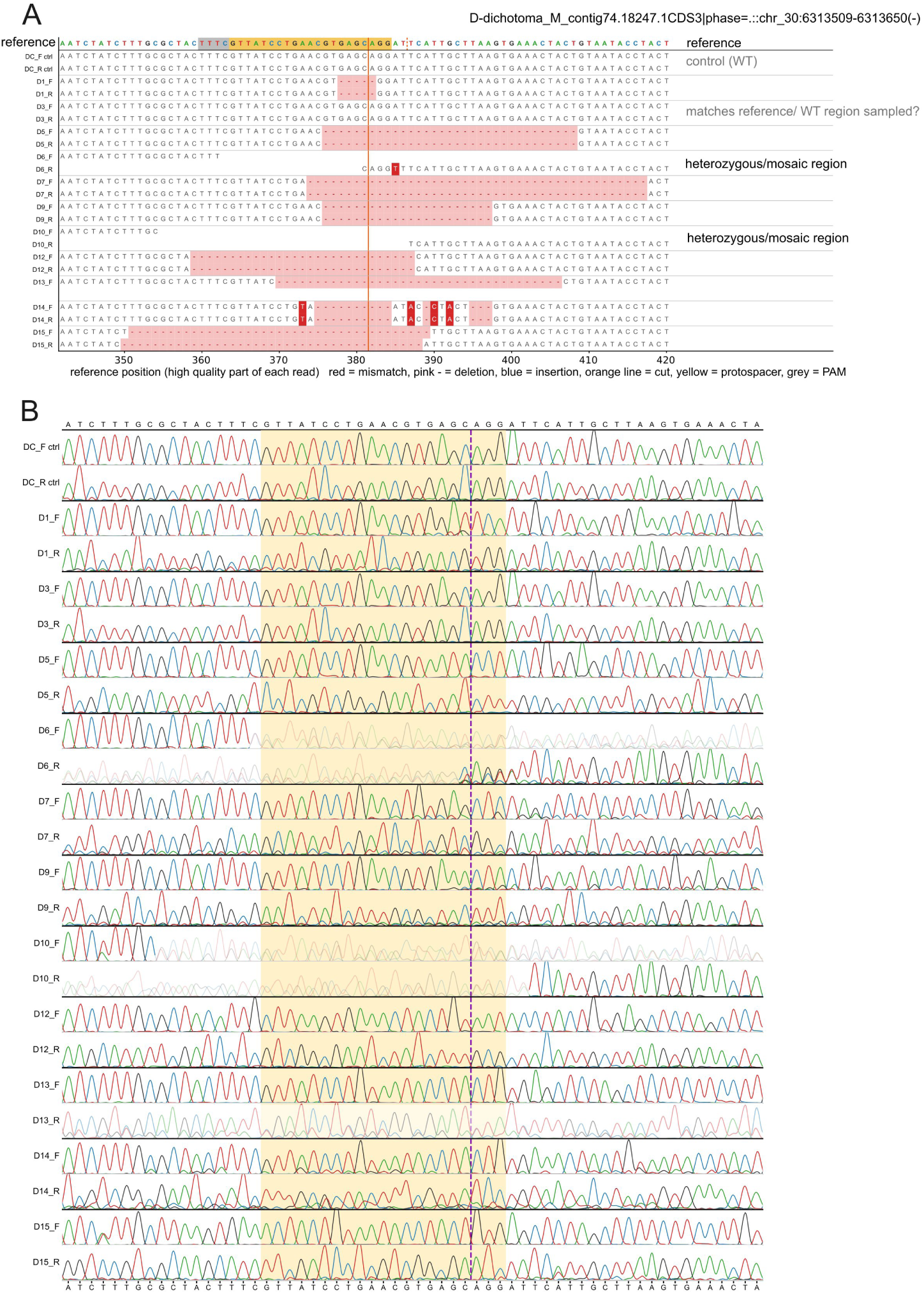
Genotyping of CRISPR/Cas12a-edited *Dictyota APT* sporophyte lines. **(A)** Sanger reads aligned to the reference amplicon around the target site in exon 1. Each row shows the high-quality part of one read (Forward or Reverse) of the line given on the left; the reference is on top and the x-axis gives the position in the reference. Red, mismatch; pink dashes, deletion; blue, insertion; yellow, protospacer; grey, PAM; orange line, predicted cut site. Blank stretches are regions without high-quality sequence; reads that break off at the cut and resume beyond it indicate mixed alleles (heterozygous or mosaic lines, labelled on the right). The untreated controls match the reference. **(B)** Chromatograms of the same reads. Each read is aligned to the reference on its own side of the cut, forward reads to the left and reverse reads to the right, and then runs on base by base, so an indel shows up as a loss of correspondence beyond the cut. Yellow, protospacer; dashed line, cut site. Faded traces mark low-quality or unanchored parts of a read. Because a mixture of alleles with different indel sizes produces two superimposed sequences downstream of the cut, lines whose forward and reverse reads both deteriorated at the cut were scored as heterozygous or mosaic rather than assigned a single allele.

**Figure S4.**
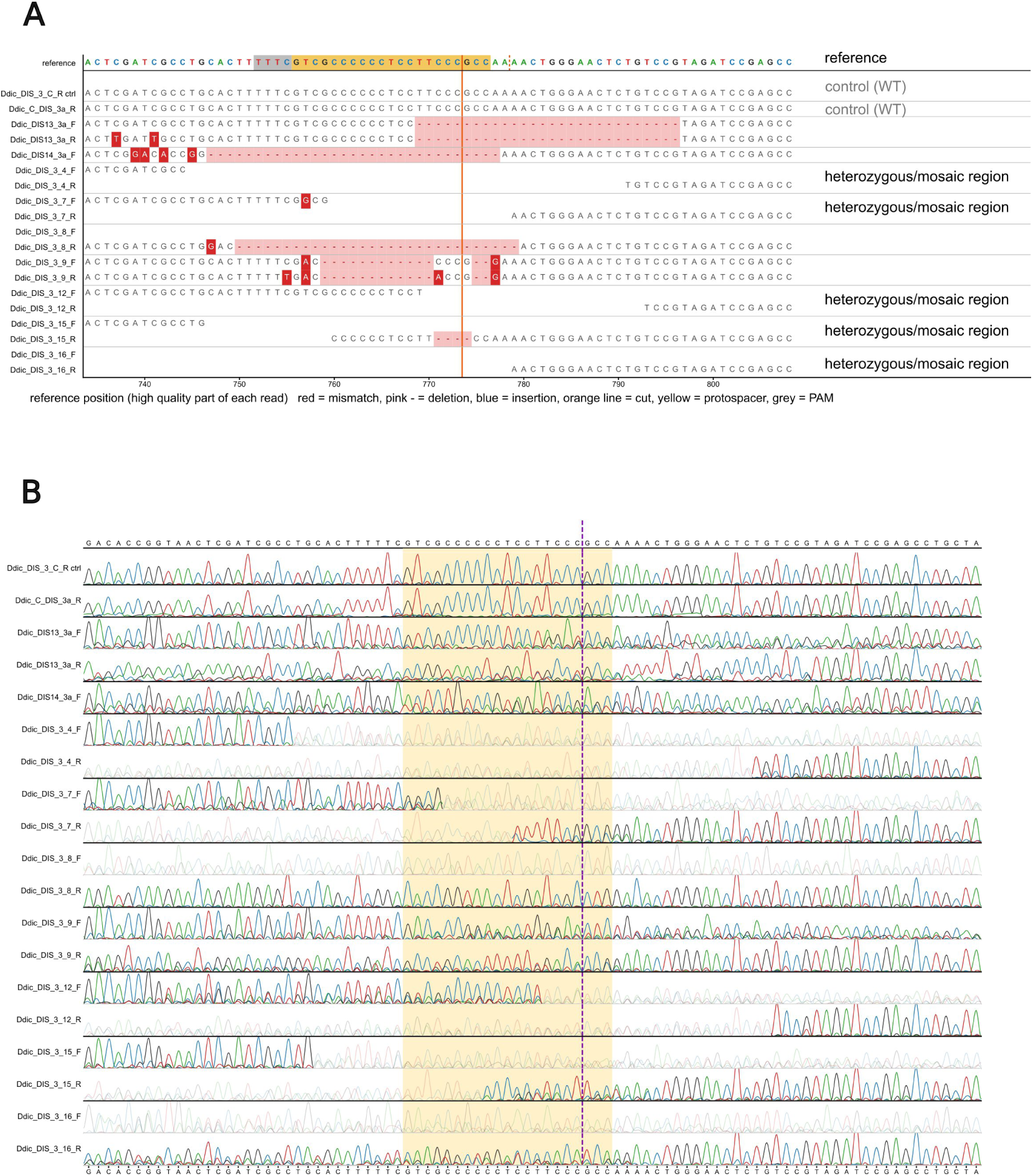
Genotyping of CRISPR/Cas12a-edited *Dictyota DIS* sporophyte lines. **(A)** Sanger reads aligned to the reference amplicon around the target site in exon 3. Each row shows the high-quality part of one read (Forward or Reverse) of the line given on the left; the reference is on top and the x-axis gives the position in the reference. Red, mismatch; pink dashes, deletion; blue, insertion; yellow, protospacer; grey, PAM; orange line, predicted cut site. Blank stretches are regions without high-quality sequence; reads that break off at the cut and resume beyond it indicate mixed alleles (heterozygous or mosaic lines, labelled on the right). The untreated controls match the reference. **(B)** Chromatograms of the same reads. Each read is aligned to the reference on its own side of the cut, forward reads to the left and reverse reads to the right, and then runs on base by base, so an indel shows up as a loss of correspondence beyond the cut. Yellow, protospacer; dashed line, cut site. Faded traces mark low-quality or unanchored parts of a read. Because a mixture of alleles with different indel sizes produces two superimposed sequences downstream of the cut, lines whose forward and reverse reads both deteriorated at the cut were scored as heterozygous or mosaic rather than assigned a single allele.

## Supplemental Tables Legends

**Table S1. TBCC-domain proteins identified in brown algae and other stramenopiles.** Predicted proteomes of 53 brown algae and 14 other stramenopiles were searched with profile HMMs of the RP2, TBCC and TBCCd1/DISTAG subfamilies, built from the reference alignment of Godfroy et al. (2017). Each hit was assigned to the subfamily whose profile gave the highest score. In total, 299 sequences were identified: 159 RP2, 64 TBCC and 76 TBCCd1/DISTAG. Proteomes without a hit for a given subfamily are not listed for that subfamily. Alternative isoforms of the same gene are listed separately.

**Table S2. crRNA and primer sequences used for Cas12a genome editing in *Dictyota*.** Top, crRNAs; bottom, primers used to amplify the target regions for genotyping. Protospacer sequences exclude the TTTV protospacer-adjacent motif. In the crRNA names, Ddic refers to Dictyota, ex indicates the targeted exon, the number indicates the position of the protospacer within the exon, and f and r indicate the forward and reverse strand respectively. The guide designations used in the main text and figures are given in the first column.

**Table S3. CRISPR/Cas12 mutagenesis of APT in *Dictyota dichotoma*.** Each row is an independent experiment. Embryos were selected on 2-fluoroadenine (2-FA), and surviving (resistant) embryos were counted as putative apt mutants. Efficiency = resistant / (resistant + dead) embryos. Dead eggs (unfertilised) died before selection and were not included.

**Table S4. CRISPR/Cas12 mutagenesis of DISTAG in *Dictyota dichotoma*.** Each row is an independent experiment, using either all five or two sgRNAs (Ddic_DIS_3_56r_Cas12_cpf; Ddic_DIS_3_101f_Cas12_cpf). Embryos were scored phenotypically as wild type (wide, light-coloured, flattened blades putative DIS/DIS), mutant (dark, with a collapsed meristem; putative dis/dis) or showing both phenotypes (putative DIS/dis). Percentages are relative to the total number of scored embryos. Total efficiency = DIS/dis + dis/dis.

**Table S5. Segregation of the dis phenotype in haploid progeny of edited diploid sporophytes.** Numbers of mutant (dis), normally developing (wt) and mosaic progeny recovered from four edited parental sporophytes (*Ddic_Dis_C, Ddic_dis_14, Ddic_dis_10*) in three independent spore releases, shown as counts (top) and as percentages of the progeny scored per release (bottom). Progeny were scored 4 weeks after release. In all parents the two main classes segregated consistent with Mendelian segregation of a single nuclear locus.

## References

Abramson, J., Adler, J., Dunger, J., Evans, R., Green, T., Pritzel, A., Ronneberger, O., Will-more, L., Ballard, A. J., Bambrick, J., et al. (2024). Accurate structure prediction of biomolecular interactions with AlphaFold 3. Nature 630, 493–500.

André, J., Harrison, S., Towers, K., Qi, X., Vaughan, S., McKean, P. G. and Ginger, M. L. (2013). The tubulin cofactor C family member TBCCD1 orchestrates cytoskeletal filament formation. J. Cell Sci. 126, 5350–5356.

Badis, Y., Scornet, D., Harada, M., Caillard, C., Godfroy, O., Raphalen, M., Gachon, C. M. M., Coelho, S. M., Motomura, T., Nagasato, C., et al. (2021). Targeted CRISPR-Cas9-based gene knockouts in the model brown alga *Ectocarpus*. New Phytol. 231, 2077–2091.

Bartolini, F., Bhamidipati, A., Thomas, S., Schwahn, U., Lewis, S. A. and Cowan, N. J. (2002). Functional Overlap between Retinitis Pigmentosa 2 Protein and the Tubulin-specific Chaperone Cofactor C. J. Biol. Chem. 277, 14629–14634.

Batista, R. A., Wang, L., Bogaert, K. A. and Coelho, S. M. (2024). Insights into the molecular bases of multicellular development from brown algae. Development 151, dev203004.

Blaby-Haas, C. E. and Merchant, S. S. (2019). Comparative and Functional Algal Genomics. Annu. Rev. Plant Biol. 70, 605–638.

Blomme, J., Liu, X., Jacobs, T. B. and De Clerck, O. (2021). A molecular toolkit for the green seaweed *Ulva mutabilis*. Plant Physiol. 186, 1442–1454.

Blomme, J., Arraiza Ribera, J., De Clerck, O. and Jacobs, T. B. (2025). Consolidating Ulva functional genomics: gene editing and new selection systems. New Phytol. 246, 1710–1723.

Bogaert, K., Beeckman, T. and De Clerck, O. (2016). Abiotic regulation of growth and fertility in the sporophyte of *Dictyota dichotoma* (Hudson) J.V. Lamouroux (Dictyotales, Phaeophyceae). J. Appl. Phycol. 28, 2915–2924.

Bogaert, K. A., Beeckman, T., and De Clerck, O. (2017a). Two-step cell polarization in algal zygotes. Nature plants, 3, 16221.

Bogaert, K. A., Beeckman, T. and De Clerck, O. (2017b). Egg activation-triggered shape change in the *Dictyota dichotoma* (Phaeophyceae) zygote is actin–myosin and secretion dependent. Ann. Bot. 120, 529–538.

Bogaert, K. A., Zakka, E. E., Coelho, S. M. and De Clerck, O. (2023). Polarization of brown algal zygotes. Semin. Cell Dev. Biol. 134, 90–102

Bogaert, K. A., Ratchinski, P., Zheng, M., Haas, F. B., Batista, R. A. and Coelho, S. M. (2026). Evolutionary flexibility in the maternal-to-zygotic transition reveals alternative routes to embryogenesis across eukaryotes. bioRxiv 10.64898/2026.04.21.719843

Bringloe, T. T., Starko, S., Wade, R. M., Vieira, C., Kawai, H., Clerck, O. D., Cock, J. M., Coelho, S. M., Destombe, C., Valero, M., et al. (2020). Phylogeny and evolution of the brown algae. Crit. Rev. Plant Sci. 39, 281–321.

Brownlee, C., Bouget, F. Y. and Corellou, F. (2001). Choosing sides: establishment of polarity in zygotes of fucoid algae. Semin. Cell Dev. Biol. 12, 345–351.

Camacho, C., Coulouris, G., Avagyan, V., Ma, N., Papadopoulos, J., Bealer, K., & Madden, T. L. (2009). BLAST+: architecture and applications. BMC bioinformatics, 10(1), 421.

Charrier, B., Coelho, S. M., Le Bail, A., Tonon, T., Michel, G., Potin, P., Kloareg, B., Boyen, C., Peters, A. F. and Cock, J. M. (2008). Development and physiology of the brown alga *Ectocarpus siliculosus*: two centuries of research. New Phytol. 177, 319–332.

Coelho, S. M. and Cock, J. M. (2020). Brown Algal Model Organisms. Annu. Rev. Genet. 54, 71–92.

Coelho, S. M. and Weijers, D. (2026). Convergent paths to multicellular complexity in photosynthetic eukaryotes. Curr. Biol. 36, 668–676.

Coelho, S. M. B., Brownlee, C. and Bothwell, J. H. F. (2008). A tip-high, Ca(2+)-interdependent, reactive oxygen species gradient is associated with polarized growth in *Fucus serratus* zygotes. Planta 227, 1037–1046.

Coelho, S. M., Peters, A. F., Müller, D. and Cock, J. M. (2020). *Ectocarpus*: an evo-devo model for the brown algae. EvoDevo 11, 19.

Concordet, J.-P. and Haeussler, M. (2018). CRISPOR: intuitive guide selection for CRISPR/Cas9 genome editing experiments and screens. Nucleic Acids Res. 46, 242–245.

Denoeud, F., Godfroy, O., Cruaud, C., Heesch, S., Nehr, Z., Tadrent, N., Couloux, A., Brillet-Guéguen, L., Delage, L., Mckeown, D., et al. (2024). Evolutionary genomics of the emergence of brown algae as key components of coastal ecosystems. Cell 187, 6943–6965.e39.

Eddy, S. R. (2011). Accelerated Profile HMM Searches. PLoS Comput. Biol. 7, e1002195.

Ewing, B. and Green, P. (1998). Base-Calling of Automated Sequencer Traces Using *Phred.* II. Error Probabilities. Genome Res. 8, 186–194.

Feldman, J. L. and Marshall, W. F. (2009). ASQ2 encodes a TBCC-like protein required for mother-daughter centriole linkage and mitotic spindle orientation. Curr. Biol. CB 19, 1238–1243.

Fritsch, F. E. (1945). The Structure and Reproduction of the Algae, Vol. II. Foreword, Phaeophyceae, Rhodophyceae, Myxophyceae. Cambridge: Cambridge University Press.

Gaillard, J., L’Hardy-Halos, M.-T. and Pellegrini, L. (1986). Morphogenese du *Dictyota dichotoma* (Huds.) Lamouroux (Phaeophyta). II. Ontogenese du thalle et cytologie ultra-structurale des differents types de cellules. Phycologia 25, 340–357.

Godfroy, O., Uji, T., Nagasato, C., Lipinska, A. P., Scornet, D., Peters, A. F., Avia, K., Colin, S., Mignerot, L., Motomura, T., et al. (2017). DISTAG/TBCCd1 Is Required for Basal Cell Fate Determination in *Ectocarpus*. Plant Cell 29, 3102–3122.

Goncalves, J., Nolasco, S., Nascimento, R., Lopez Fanarraga, M., Zabala, J. C. and Soares, H. (2010). TBCCD1, a new centrosomal protein, is required for centrosome and Golgi apparatus positioning. EMBO Rep. 11, 194–200.

Ichihara, K., Yamazaki, T. and Kawano, S. (2022). Genome editing using a DNA-free clustered regularly interspaced short palindromic repeats-CAS9 system in green seaweed *Ulva prolifera*. Phycol. Res. 70, 50–56.

Ichihara, K., Yamazaki, T. and Kawano, S. (2024). Cas9 ribonucleoproteins and single-strand oligodeoxynucleotides enable targeted insertional mutagenesis in Ulva. Algal Res. 81, 103598.

Katsaros, C. and Galatis, B. (1985). Ultrastructural studies on thallus development in *Dictyota dichotoma* (Phaeophyta, Dictyotales). Br. Phycol. J. 20, 263–276.

Kumke, J. (1973). Beiträge zur periodizität der Oogon-Entleerung bei *Dictyota dichotoma* (Phaeophyta). Z. Pflanzenphysiol. 70, 191–210.

Liu, P., Vigneau, J., Craig, R. J., Barrera-Redondo, J., Avdievich, E., Martinho, C., Borg, M., Haas, F. B., Liu, C. and Coelho, S. M. (2024). 3D chromatin maps of a brown alga reveal U/V sex chromosome spatial organization. Nat. Commun. 15, 9590.

Lotharukpong, J. S., Zheng, M., Luthringer, R., Liesner, D., Drost, H.-G. and Coelho, S. M. (2024). A transcriptomic hourglass in brown algae. Nature 635, 129–135.

Martinho, C., Hoshino, M., Raphalen, M., Bukhanets, V., Kerur, A., Bogaert, K. A., Luthringer, R. and Coelho, S. M. (2026). Efficient CRISPR-Cas genome editing in brown algae. Cell Rep. Methods 101273.

Paix, A., Raphalen, M., Avdievich, E., Agullo, F., Luthringer, R. and Coelho, S. M. (2026). Endogenous gene tagging in the model brown alga *Ectocarpus* using a simplified CRISPR/Cas method. bioRxiv 10.64898/2026.08.21.745938

Pertea, G. and Pertea, M. (2020). GFF Utilities: GffRead and GffCompare. F1000Research 9, 304.

Shen, Y., Motomura, T., Ichihara, K., Matsuda, Y., Yoshimura, K., Kosugi, C. and Nagasato, C. (2023). Application of CRISPR-Cas9 genome editing by microinjection of gametophytes of Saccharina japonica (Laminariales, Phaeophyceae). J. Appl. Phycol. 35, 1431–1441.

Steinegger, M., Meier, M., Mirdita, M., Vohringer, H., Haunsberger, S. J. and Soding, J. (2019). HH-suite3 for fast remote homology detection and deep protein annotation. BMC Bioinformatics 20, 473.

Veltel, S., Gasper, R., Eisenacher, E. and Wittinghofer, A. (2008). The retinitis pigmentosa 2 gene product is a GTPase-activating protein for Arf-like 3. Nat. Struct. Mol. Biol. 15, 373–380.

Vigneau, J., Lotharukpong, J. S., Liu, P., Luthringer, R., Lombard, B., Loew, D., Haas, F. B., Borg, M. and Coelho, S. M. (2026). Evolution of a distinct chromatin regulatory landscape in brown algae. Nat. Ecol. Evol. 10, 779–793.

Yang, Z. (2008). Cell Polarity Signaling in *Arabidopsis*. Annu. Rev. Cell Dev. Biol. 24, 551–575.

Zimmermann, L., Stephens, A., Nam, S.-Z., Rau, D., Kubler, J., Lozajic, M., Gabler, F., Soding, J., Lupas, A. N. and Alva, V. (2018). A Completely Reimplemented MPI Bioinformatics Toolkit with a New HHpred Server at its Core. J. Mol. Biol. 430, 2237–2243.

