## Supplemental Tables S1-S5 for "Conserved control of cell differentiation across divergent brown algal body plans revealed by diploid genome editing"

**Table S1.** TBCC-domain proteins identified in brown algae and other stramenopiles. Predicted proteomes of 53 brown algae and 14 other stramenopiles were searched with profile HMMs of the RP2, TBCC and TBCCd1/DISTAG subfamilies, built from the reference alignment of Godfroy et al. (2017). Each hit was assigned to the subfamily whose profile gave the highest score. In total, 299 sequences were identified: 159 RP2, 64 TBCC and 76 TBCCd1/DISTAG. Proteomes without a hit for a given subfamily are not listed for that subfamily. Alternative isoforms of the same gene are listed separately.

| sequence | Species | subfamily | score | evalue |
|---|---|---|---|---|
| Ascophyllum_nodosum\|prot_A-nodosum_M_contig1361.4.26 | Ascophyllum_nodosum | RP2 | 173.7 | 2.10E-55 |
| Ascophyllum_nodosum\|prot_A-nodosum_M_contig181.38.1 | Ascophyllum_nodosum | RP2 | 37.6 | 3.00E-13 |
| Ascophyllum_nodosum\|prot_A-nodosum_M_contig516.43.2 | Ascophyllum_nodosum | RP2 | 174.4 | 1.30E-55 |
| Ascophyllum_nodosum\|prot_A-nodosum_M_contig409.17.2 | Ascophyllum_nodosum | TBCC | 145.1 | 1.70E-46 |
| Ascophyllum_nodosum\|prot_A-nodosum_M_contig2041.8.1 | Ascophyllum_nodosum | TBCCd1 | 165.9 | 4.50E-53 |
| Asterocladon_lobatum\|Alob_scaffold_02145.g7444.t2 | Asterocladon_lobatum | RP2 | 37.3 | 3.80E-13 |
| Asterocladon_lobatum\|Alob_scaffold_01746.g6645.t1 | Asterocladon_lobatum | TBCC | 164.3 | 2.00E-52 |
| Asterocladon_lobatum\|Alob_scaffold_03459.g9356.t1 | Asterocladon_lobatum | TBCCd1 | 85.3 | 4.00E-28 |
| Aureococcus_anophagefferens\|EGB07258 | Aureococcus_anophagefferens | RP2 | 152.4 | 8.30E-49 |
| Aureococcus_anophagefferens\|EGB12705 | Aureococcus_anophagefferens | RP2 | 85.6 | 4.10E-28 |
| Chordaria_linearis\|prot_C-linearis_contig19.4627.1 | Chordaria_linearis | RP2 | 28.3 | 2.30E-10 |
| Chordaria_linearis\|prot_C-linearis_contig2.4794.1 | Chordaria_linearis | RP2 | 189.2 | 3.30E-60 |
| Chordaria_linearis\|prot_C-linearis_contig63.13630.1 | Chordaria_linearis | RP2 | 188 | 7.50E-60 |
| Chordaria_linearis\|prot_C-linearis_contig18.4360.1 | Chordaria_linearis | TBCC | 183.9 | 1.70E-58 |
| Choristocarpus_tenellus\|prot_C-tenellus_contig12488.1.1 | Choristocarpus_tenellus | RP2 | 142.6 | 9.10E-46 |
| Choristocarpus_tenellus\|prot_C-tenellus_contig9245.3.1 | Choristocarpus_tenellus | RP2 | 172.6 | 4.40E-55 |
| Chrysoparadoxa_australica\|prot_C-australica_Contig_98.14.2 | Chrysoparadoxa_australica | RP2 | 162.9 | 4.50E-52 |
| Chrysoparadoxa_australica\|prot_C-australica_Contig_475.11.2 | Chrysoparadoxa_australica | TBCC | 156.5 | 5.00E-50 |
| Chrysoparadoxa_australica\|prot_C-australica_Contig_146.11.6 | Chrysoparadoxa_australica | TBCCd1 | 150.9 | 1.90E-48 |
| Cladosiphon_okamuranus\|Cok_S_g13654.t1 | Cladosiphon_okamuranus | RP2 | 182.5 | 3.80E-58 |
| Cladosiphon_okamuranus\|Cok_S_g7714.t1 | Cladosiphon_okamuranus | RP2 | 184.9 | 7.10E-59 |
| Cladosiphon_okamuranus\|Cok_S_g8035.t1 | Cladosiphon_okamuranus | RP2 | 26.4 | 8.80E-10 |
| Cladosiphon_okamuranus\|Cok_S_g10102.t1 | Cladosiphon_okamuranus | TBCC | 175.3 | 7.90E-56 |
| Cladosiphon_okamuranus\|Cok_S_g7486.t1 | Cladosiphon_okamuranus | TBCCd1 | 178.8 | 4.50E-57 |
| Desmarestia_dudresnayi\|Ddudresnayi_g12491.t1 | Desmarestia_dudresnayi | RP2 | 182.8 | 3.00E-58 |
| Desmarestia_dudresnayi\|Ddudresnayi_g27700.t1 | Desmarestia_dudresnayi | RP2 | 184.3 | 1.10E-58 |
| Desmarestia_dudresnayi\|Ddudresnayi_g6974.t1 | Desmarestia_dudresnayi | TBCC | 178.3 | 9.00E-57 |
| Desmarestia_dudresnayi\|Ddudresnayi_g9370.t1 | Desmarestia_dudresnayi | TBCCd1 | 177.4 | 1.20E-56 |
| Desmarestia_dudresnayi\|Ddudresnayi_g9371.t1 | Desmarestia_dudresnayi | TBCCd1 | 177.2 | 1.40E-56 |
| Desmarestia_herbacea\|mRNA_D-herbacea_M_contig40.10279.1 | Desmarestia_herbacea | RP2 | 182.9 | 2.90E-58 |
| Desmarestia_herbacea\|mRNA_D-herbacea_M_contig110.998.1 | Desmarestia_herbacea | TBCC | 181.8 | 7.30E-58 |
| Desmarestia_herbacea\|mRNA_D-herbacea_M_contig68.13962.1 | Desmarestia_herbacea | TBCCd1 | 177.1 | 1.40E-56 |
| Dictyota_dichotoma\|prot_D-dichotoma_M_contig386.12784.1 | Dictyota_dichotoma | RP2 | 181.5 | 8.10E-58 |
| Dictyota_dichotoma\|prot_D-dichotoma_M_contig70.17897.1 | Dictyota_dichotoma | RP2 | 179.5 | 3.30E-57 |
| Dictyota_dichotoma\|prot_D-dichotoma_M_contig512.15537.1 | Dictyota_dichotoma | TBCC | 160.3 | 3.40E-51 |
| Dictyota_dichotoma\|prot_D-dichotoma_M_contig74.18247.1 | Dictyota_dichotoma | TBCCd1 | 171.6 | 7.30E-55 |
| Discosporangium_mesarthrocarpum\|prot_D-mesarthrocarpus_Contig2720.1.1 | Discosporangium_mesarthrocarpum | RP2 | 177.2 | 1.70E-56 |
| Discosporangium_mesarthrocarpum\|prot_D-mesarthrocarpus_Contig8342.1.1 | Discosporangium_mesarthrocarpum | TBCCd1 | 161.7 | 8.60E-52 |
| Ectocarpus_Ec05\|Ec-15_001460.1 | Ectocarpus_Ec05 | RP2 | 22.3 | 0.000000016 |
| Ectocarpus_Ec05\|Ec-19_004040.1 | Ectocarpus_Ec05 | RP2 | 188.5 | 5.40E-60 |
| Ectocarpus_Ec05\|Ec-21_002520.1 | Ectocarpus_Ec05 | RP2 | 193.8 | 1.20E-61 |

| | | | | |
|---|---|---|---|---|
| Ectocarpus_Ec05\|Ec-08_006490.1 | Ectocarpus_Ec05 | TBCC | 179.9 | 2.80E-57 |
| Ectocarpus_Ec05\|Ec-05_001860.1 | Ectocarpus_Ec05 | TBCCd1 | 180.7 | 1.20E-57 |
| Ectocarpus_Ec25\|Ec-15_001460.1 | Ectocarpus_Ec25 | RP2 | 19.8 | 0.000000094 |
| Ectocarpus_Ec25\|Ec-19_004040.1 | Ectocarpus_Ec25 | RP2 | 189.5 | 2.60E-60 |
| Ectocarpus_Ec25\|Ec-21_002520.1 | Ectocarpus_Ec25 | RP2 | 194.6 | 6.90E-62 |
| Ectocarpus_Ec25\|Ec-08_006490.1 | Ectocarpus_Ec25 | TBCC | 183.9 | 1.60E-58 |
| Ectocarpus_Ec25\|Ec-05_001860.1 | Ectocarpus_Ec25 | TBCCd1 | 180.6 | 1.20E-57 |
| Ectocarpus_Ec32\|Ec-15_001460.1 | Ectocarpus_Ec32 | RP2 | 19.8 | 0.000000094 |
| Ectocarpus_Ec32\|Ec-19_004040.1 | Ectocarpus_Ec32 | RP2 | 190.4 | 1.40E-60 |
| Ectocarpus_Ec32\|Ec-21_002520.1 | Ectocarpus_Ec32 | RP2 | 193.7 | 1.30E-61 |
| Ectocarpus_Ec32\|Ec-08_006490.1 | Ectocarpus_Ec32 | TBCC | 183.9 | 1.60E-58 |
| Ectocarpus_Ec32\|Ec-05_001860.1 | Ectocarpus_Ec32 | TBCCd1 | 180.6 | 1.20E-57 |
| Ectocarpus_crouaniorum\|prot_E-crouaniorum-2c_M_contig19.4531.1 | Ectocarpus_crouaniorum | RP2 | 178.2 | 8.20E-57 |
| Ectocarpus_crouaniorum\|prot_E-crouaniorum-2c_M_contig20.5866.1 | Ectocarpus_crouaniorum | RP2 | 18.9 | 0.00000018 |
| Ectocarpus_crouaniorum\|prot_E-crouaniorum-2c_M_contig22.6586.1 | Ectocarpus_crouaniorum | RP2 | 186.1 | 3.00E-59 |
| Ectocarpus_crouaniorum\|prot_E-crouaniorum-2c_M_contig3.8735.1 | Ectocarpus_crouaniorum | TBCC | 182 | 6.60E-58 |
| Ectocarpus_crouaniorum\|prot_E-crouaniorum-2c_M_contig56.14979.1 | Ectocarpus_crouaniorum | TBCCd1 | 178.2 | 6.70E-57 |
| Ectocarpus_fasciculatus\|prot_E-fasciculatus_M_contig5.14071.1 | Ectocarpus_fasciculatus | RP2 | 22.7 | 0.000000012 |
| Ectocarpus_fasciculatus\|prot_E-fasciculatus_M_contig61.15630.1 | Ectocarpus_fasciculatus | RP2 | 192.6 | 2.80E-61 |
| Ectocarpus_fasciculatus\|prot_E-fasciculatus_M_contig73.17025.1 | Ectocarpus_fasciculatus | RP2 | 174.3 | 1.30E-55 |
| Ectocarpus_fasciculatus\|prot_E-fasciculatus_M_contig160.4564.1 | Ectocarpus_fasciculatus | TBCC | 179.2 | 4.80E-57 |
| Ectocarpus_fasciculatus\|prot_E-fasciculatus_M_contig74.17128.1 | Ectocarpus_fasciculatus | TBCCd1 | 182.3 | 3.70E-58 |
| Ectocarpus_siliculosus\|prot_E-siliculosus-1a_M_contig37.10274.1 | Ectocarpus_siliculosus | RP2 | 124.6 | 3.30E-40 |
| Ectocarpus_siliculosus\|prot_E-siliculosus-1a_M_contig37.10275.1 | Ectocarpus_siliculosus | RP2 | 52.8 | 5.90E-18 |
| Ectocarpus_siliculosus\|prot_E-siliculosus-1a_M_contig83.16450.1 | Ectocarpus_siliculosus | RP2 | 190.3 | 1.50E-60 |
| Ectocarpus_siliculosus\|prot_E-siliculosus-1a_M_contig84.16537.1 | Ectocarpus_siliculosus | RP2 | 22.4 | 0.000000015 |
| Ectocarpus_siliculosus\|prot_E-siliculosus-1a_M_contig300.9238.1 | Ectocarpus_siliculosus | TBCC | 178.2 | 9.50E-57 |
| Ectocarpus_siliculosus\|prot_E-siliculosus-1a_M_contig234.7273.1 | Ectocarpus_siliculosus | TBCCd1 | 180.6 | 1.20E-57 |
| Ectocarpus_sp1\|prot_Ecto-sp1_M_contig1690.3720.1 | Ectocarpus_sp1 | RP2 | 109.9 | 1.20E-35 |
| Ectocarpus_sp1\|prot_Ecto-sp1_M_contig2978.8010.1 | Ectocarpus_sp1 | RP2 | 22.2 | 0.000000017 |
| Ectocarpus_sp1\|prot_Ecto-sp1_M_contig320.8601.1 | Ectocarpus_sp1 | RP2 | 170.7 | 1.70E-54 |
| Ectocarpus_sp1\|prot_Ecto-sp1_M_contig3908.10242.1 | Ectocarpus_sp1 | TBCC | 180.1 | 2.40E-57 |
| Ectocarpus_sp1\|prot_Ecto-sp1_M_contig478.11948.1 | Ectocarpus_sp1 | TBCCd1 | 180 | 1.90E-57 |
| Ectocarpus_sp12\|prot_Ecto-sp12_S_contig10858.1210.1 | Ectocarpus_sp12 | RP2 | 166.9 | 2.60E-53 |
| Ectocarpus_sp12\|prot_Ecto-sp12_S_contig1676.8021.1 | Ectocarpus_sp12 | RP2 | 108.3 | 3.70E-35 |
| Ectocarpus_sp12\|prot_Ecto-sp12_S_contig19273.10409.1 | Ectocarpus_sp12 | RP2 | 22.9 | 0.00000001 |
| Ectocarpus_sp12\|prot_Ecto-sp12_S_contig2214.12923.1 | Ectocarpus_sp12 | RP2 | 170.4 | 2.20E-54 |
| Ectocarpus_sp12\|prot_Ecto-sp12_S_contig975.47983.1 | Ectocarpus_sp12 | RP2 | 187.7 | 9.60E-60 |
| Ectocarpus_sp12\|prot_Ecto-sp12_S_contig23642.14053.1 | Ectocarpus_sp12 | TBCC | 184.4 | 1.20E-58 |
| Ectocarpus_sp12\|prot_Ecto-sp12_S_contig71330.38662.1 | Ectocarpus_sp12 | TBCCd1 | 149.9 | 3.80E-48 |
| Ectocarpus_sp13\|prot_Ecto-sp13_S_contig1024.267.1 | Ectocarpus_sp13 | RP2 | 175 | 8.20E-56 |
| Ectocarpus_sp13\|prot_Ecto-sp13_S_contig3786.12323.1 | Ectocarpus_sp13 | RP2 | 112.4 | 1.90E-36 |
| Ectocarpus_sp13\|prot_Ecto-sp13_S_contig13240.2612.1 | Ectocarpus_sp13 | TBCC | 156 | 7.30E-50 |
| Ectocarpus_sp13\|prot_Ecto-sp13_S_contig7274.18422.1 | Ectocarpus_sp13 | TBCCd1 | 112.8 | 1.20E-36 |
| Ectocarpus_sp2\|prot_Ecto-sp2_M_contig1739.4486.1 | Ectocarpus_sp2 | RP2 | 22.3 | 0.000000016 |
| Ectocarpus_sp2\|prot_Ecto-sp2_M_contig4393.10498.1 | Ectocarpus_sp2 | RP2 | 114.8 | 3.50E-37 |
| Ectocarpus_sp2\|prot_Ecto-sp2_M_contig6527.13444.1 | Ectocarpus_sp2 | RP2 | 197.1 | 1.20E-62 |
| Ectocarpus_sp2\|prot_Ecto-sp2_M_contig3728.9454.1 | Ectocarpus_sp2 | TBCC | 157.4 | 2.60E-50 |
| Ectocarpus_sp2\|prot_Ecto-sp2_M_contig21081.5852.1 | Ectocarpus_sp2 | TBCCd1 | 72.6 | 3.50E-24 |
| Ectocarpus_sp3\|prot_Ecto-sp3_M_contig591.13765.1 | Ectocarpus_sp3 | RP2 | 141.3 | 2.20E-45 |
| Ectocarpus_sp3\|prot_Ecto-sp3_M_contig825.16471.1 | Ectocarpus_sp3 | RP2 | 172.2 | 6.20E-55 |

| | | | | |
|---|---|---|---|---|
| Ectocarpus_sp3\|prot_Ecto-sp3_M_contig132.2489.1 | Ectocarpus_sp3 | TBCC | 179.2 | 4.90E-57 |
| Ectocarpus_sp5\|prot_Ecto-sp5_M_contig15881.4305.1 | Ectocarpus_sp5 | RP2 | 35.2 | 1.70E-12 |
| Ectocarpus_sp5\|prot_Ecto-sp5_M_contig16103.4455.1 | Ectocarpus_sp5 | RP2 | 141.9 | 1.40E-45 |
| Ectocarpus_sp5\|prot_Ecto-sp5_M_contig23324.8211.1 | Ectocarpus_sp5 | RP2 | 124.7 | 3.00E-40 |
| Ectocarpus_sp5\|prot_Ecto-sp5_M_contig44060.14456.1 | Ectocarpus_sp5 | TBCCd1 | 62.3 | 5.30E-21 |
| Ectocarpus_sp6\|prot_Ecto-sp6_S_contig40.11810.1 | Ectocarpus_sp6 | RP2 | 170.5 | 2.00E-54 |
| Ectocarpus_sp6\|prot_Ecto-sp6_S_contig72.16381.1 | Ectocarpus_sp6 | RP2 | 21.4 | 0.000000031 |
| Ectocarpus_sp6\|prot_Ecto-sp6_S_contig8.17113.1 | Ectocarpus_sp6 | RP2 | 190.3 | 1.50E-60 |
| Ectocarpus_sp6\|prot_Ecto-sp6_S_contig144.3808.1 | Ectocarpus_sp6 | TBCC | 184.2 | 1.40E-58 |
| Ectocarpus_sp6\|prot_Ecto-sp6_S_contig169.4770.1 | Ectocarpus_sp6 | TBCC | 181.5 | 9.30E-58 |
| Ectocarpus_sp6\|prot_Ecto-sp6_S_contig172.5014.1 | Ectocarpus_sp6 | TBCCd1 | 180.6 | 1.20E-57 |
| Ectocarpus_sp8\|prot_Ecto-sp8_S_contig11229.1588.1 | Ectocarpus_sp8 | RP2 | 145.2 | 1.40E-46 |
| Ectocarpus_sp8\|prot_Ecto-sp8_S_contig2493.11596.1 | Ectocarpus_sp8 | RP2 | 139.6 | 7.40E-45 |
| Ectocarpus_sp8\|prot_Ecto-sp8_S_contig27194.12819.1 | Ectocarpus_sp8 | RP2 | 21 | 0.000000042 |
| Ectocarpus_sp8\|prot_Ecto-sp8_S_contig39817.18125.1 | Ectocarpus_sp8 | RP2 | 20.2 | 0.000000075 |
| Ectocarpus_sp8\|prot_Ecto-sp8_S_contig88901.30019.1 | Ectocarpus_sp8 | RP2 | 53.5 | 3.40E-18 |
| Ectocarpus_sp8\|prot_Ecto-sp8_S_contig3345.15721.1 | Ectocarpus_sp8 | TBCC | 180.1 | 2.50E-57 |
| Ectocarpus_sp8\|prot_Ecto-sp8_S_contig7919.28280.1 | Ectocarpus_sp8 | TBCC | 146 | 8.90E-47 |
| Ectocarpus_sp8\|prot_Ecto-sp8_S_contig471.20542.1 | Ectocarpus_sp8 | TBCCd1 | 180.6 | 1.20E-57 |
| Ectocarpus_sp9\|prot_Ecto-sp9_F_contig1676.4200.1 | Ectocarpus_sp9 | RP2 | 189.9 | 1.90E-60 |
| Ectocarpus_sp9\|prot_Ecto-sp9_F_contig2481.6940.1 | Ectocarpus_sp9 | RP2 | 21.1 | 0.000000037 |
| Ectocarpus_sp9\|prot_Ecto-sp9_F_contig292.8059.1 | Ectocarpus_sp9 | RP2 | 168.9 | 6.30E-54 |
| Ectocarpus_sp9\|prot_Ecto-sp9_F_contig481.11781.1 | Ectocarpus_sp9 | TBCC | 181.5 | 9.30E-58 |
| Ectocarpus_subulatus\|EsuBft1158_3 | Ectocarpus_subulatus | RP2 | 155.7 | 7.90E-50 |
| Ectocarpus_subulatus\|EsuBft330_12 | Ectocarpus_subulatus | RP2 | 139.6 | 7.60E-45 |
| Ectocarpus_subulatus\|EsuBft97_15 | Ectocarpus_subulatus | RP2 | 21.5 | 0.000000028 |
| Ectocarpus_subulatus\|EsuBft1602_3 | Ectocarpus_subulatus | TBCC | 181.3 | 1.10E-57 |
| Ectocarpus_subulatus\|EsuBft300_4 | Ectocarpus_subulatus | TBCCd1 | 182.3 | 3.70E-58 |
| Feldmannia_mitchelliae\|prot_F-mitchelliae_contig1188.1390.1 | Feldmannia_mitchelliae | RP2 | 181.8 | 6.30E-58 |
| Feldmannia_mitchelliae\|prot_F-mitchelliae_contig204.5191.1 | Feldmannia_mitchelliae | RP2 | 182.7 | 3.50E-58 |
| Feldmannia_mitchelliae\|prot_F-mitchelliae_contig4133.10508.1 | Feldmannia_mitchelliae | TBCC | 169 | 6.90E-54 |
| Feldmannia_mitchelliae\|prot_F-mitchelliae_contig2568.6891.1 | Feldmannia_mitchelliae | TBCCd1 | 175.9 | 3.60E-56 |
| Fucus_distichus\|prot_F-distichus_contig19881.6572.1 | Fucus_distichus | RP2 | 31.2 | 2.80E-11 |
| Fucus_distichus\|prot_F-distichus_contig6774.15973.1 | Fucus_distichus | RP2 | 174.9 | 8.60E-56 |
| Fucus_distichus\|prot_F-distichus_contig53060.13842.1 | Fucus_distichus | TBCC | 160.6 | 2.70E-51 |
| Fucus_distichus\|prot_F-distichus_contig40962.11957.1 | Fucus_distichus | TBCCd1 | 89.5 | 2.10E-29 |
| Fucus_serratus\|prot_F-serratus_M_contig40.12445.1 | Fucus_serratus | RP2 | 32.4 | 1.20E-11 |
| Fucus_serratus\|prot_F-serratus_M_contig5563.15835.1 | Fucus_serratus | RP2 | 175 | 8.30E-56 |
| Fucus_serratus\|prot_F-serratus_M_contig601.16658.1 | Fucus_serratus | RP2 | 175 | 8.20E-56 |
| Fucus_serratus\|prot_F-serratus_M_contig1435.3114.1 | Fucus_serratus | TBCC | 159 | 8.30E-51 |
| Fucus_serratus\|prot_F-serratus_M_contig924.20641.1 | Fucus_serratus | TBCCd1 | 165.1 | 8.00E-53 |
| Halopteris_paniculata\|prot_H-paniculata_contig1723.4030.1 | Halopteris_paniculata | RP2 | 178.9 | 5.10E-57 |
| Halopteris_paniculata\|prot_H-paniculata_contig668.14566.1 | Halopteris_paniculata | RP2 | 184.3 | 1.10E-58 |
| Halopteris_paniculata\|prot_H-paniculata_contig443.11195.1 | Halopteris_paniculata | TBCC | 161 | 2.00E-51 |
| Halopteris_paniculata\|prot_H-paniculata_contig5401.12838.1 | Halopteris_paniculata | TBCCd1 | 175.9 | 3.50E-56 |
| Hapterophycus_canaliculatus\|prot_H-canaliculatus_M_contigs20210.1.1 | Hapterophycus_canaliculatus | RP2 | 108.2 | 4.00E-35 |
| Hapterophycus_canaliculatus\|prot_H-canaliculatus_M_contigs6024.1.1 | Hapterophycus_canaliculatus | RP2 | 100.2 | 1.20E-32 |
| Hapterophycus_canaliculatus\|prot_H-canaliculatus_M_contigs16191.1.1 | Hapterophycus_canaliculatus | TBCC | 123.1 | 1.10E-39 |
| Hapterophycus_canaliculatus\|prot_H-canaliculatus_M_contigs27747.1.1 | Hapterophycus_canaliculatus | TBCCd1 | 154.6 | 1.40E-49 |
| Heribaudiella_fluviatilis\|prot_H-fluviatilis_contig12903.4665.1 | Heribaudiella_fluviatilis | RP2 | 120.7 | 5.50E-39 |
| Heribaudiella_fluviatilis\|prot_H-fluviatilis_contig3777.22611.1 | Heribaudiella_fluviatilis | RP2 | 177.5 | 1.40E-56 |
| Heribaudiella_fluviatilis\|prot_H-fluviatilis_contig42370.24861.1 | Heribaudiella_fluviatilis | RP2 | 109.1 | 2.10E-35 |

| | | | | |
|---|---|---|---|---|
| Heribaudiella_fluviatilis\|prot_H-fluviatilis_contig6823.34253.1 | Heribaudiella_fluviatilis | RP2 | 135.3 | 1.60E-43 |
| Heribaudiella_fluviatilis\|prot_H-fluviatilis_contig137.5490.1 | Heribaudiella_fluviatilis | TBCC | 170.1 | 3.00E-54 |
| Heribaudiella_fluviatilis\|prot_H-fluviatilis_contig18380.9614.1 | Heribaudiella_fluviatilis | TBCC | 122.7 | 1.50E-39 |
| Heribaudiella_fluviatilis\|prot_H-fluviatilis_contig105860.1142.1 | Heribaudiella_fluviatilis | TBCCd1 | 62.7 | 4.00E-21 |
| Heribaudiella_fluviatilis\|prot_H-fluviatilis_contig25535.15185.1 | Heribaudiella_fluviatilis | TBCCd1 | 85.5 | 3.50E-28 |
| Heribaudiella_fluviatilis\|prot_H-fluviatilis_contig641.33040.1 | Heribaudiella_fluviatilis | TBCCd1 | 111.4 | 3.30E-36 |
| Heribaudiella_fluviatilis\|prot_H-fluviatilis_contig93903.40176.1 | Heribaudiella_fluviatilis | TBCCd1 | 73.6 | 1.70E-24 |
| Heterosigma_akashiwo\|prot_H-akashiwo_Contig2655.1.1 | Heterosigma_akashiwo | RP2 | 131.5 | 2.40E-42 |
| Heterosigma_akashiwo\|prot_H-akashiwo_Contig100.27.1 | Heterosigma_akashiwo | TBCCd1 | 134.1 | 3.20E-43 |
| Himanthalia_elongata\|prot_H-elongata_contig17528.4566.1 | Himanthalia_elongata | RP2 | 181.6 | 7.20E-58 |
| Himanthalia_elongata\|prot_H-elongata_contig69290.14575.1 | Himanthalia_elongata | RP2 | 107.6 | 6.20E-35 |
| Himanthalia_elongata\|prot_H-elongata_contig29467.8753.1 | Himanthalia_elongata | TBCC | 159 | 8.60E-51 |
| Himanthalia_elongata\|prot_H-elongata_contig186892.5088.1 | Himanthalia_elongata | TBCCd1 | 59.4 | 4.10E-20 |
| Himanthalia_elongata\|prot_H-elongata_contig187816.5131.1 | Himanthalia_elongata | TBCCd1 | 88.7 | 3.50E-29 |
| Laminaria_digitata\|prot_L-digitata_M_contig19671.1.1 | Laminaria_digitata | RP2 | 185.3 | 5.40E-59 |
| Laminaria_digitata\|prot_L-digitata_M_contig27793.1.1 | Laminaria_digitata | RP2 | 106.6 | 1.30E-34 |
| Laminaria_digitata\|prot_L-digitata_M_contig41903.1.1 | Laminaria_digitata | RP2 | 37.6 | 3.00E-13 |
| Laminaria_digitata\|prot_L-digitata_M_contig5592.1.1 | Laminaria_digitata | TBCC | 148.5 | 1.50E-47 |
| Laminaria_digitata\|prot_L-digitata_M_contig30685.1.1 | Laminaria_digitata | TBCCd1 | 156.5 | 3.50E-50 |
| Laminarionema_elsbetiae\|prot_L-elsbetiae_contig10586.562.1 | Laminarionema_elsbetiae | RP2 | 185.2 | 5.60E-59 |
| Laminarionema_elsbetiae\|prot_L-elsbetiae_contig8741.17772.1 | Laminarionema_elsbetiae | RP2 | 21.4 | 0.00000003 |
| Laminarionema_elsbetiae\|prot_L-elsbetiae_contig935.18348.1 | Laminarionema_elsbetiae | RP2 | 187.5 | 1.10E-59 |
| Laminarionema_elsbetiae\|prot_L-elsbetiae_contig8223.17228.1 | Laminarionema_elsbetiae | TBCC | 170.8 | 1.90E-54 |
| Laminarionema_elsbetiae\|prot_L-elsbetiae_contig32525.9677.1 | Laminarionema_elsbetiae | TBCCd1 | 63.3 | 2.50E-21 |
| Laminarionema_elsbetiae\|prot_L-elsbetiae_contig9140.18169.1 | Laminarionema_elsbetiae | TBCCd1 | 56.3 | 3.90E-19 |
| Macrocystis_pyrifera\|jgi\|Macpyr2\|8865505\|g1695.t1 | Macrocystis_pyrifera | RP2 | 181.4 | 8.40E-58 |
| Macrocystis_pyrifera\|jgi\|Macpyr2\|9509197\|fgenesh2_kg.10_#_10479_#_TRINITY_DN20561_c0_g2_i2 | Macrocystis_pyrifera | RP2 | 160.4 | 2.70E-51 |
| Macrocystis_pyrifera\|jgi\|Macpyr2\|9805612\|estExt_fgenesh2_pm.C_10221 | Macrocystis_pyrifera | TBCC | 177.1 | 2.10E-56 |
| Macrocystis_pyrifera\|jgi\|Macpyr2\|9708973\|fgenesh2_kg.29_#_2764_#_TRINITY_DN107102_c0_g1_i2 | Macrocystis_pyrifera | TBCCd1 | 111.2 | 3.70E-36 |
| Myriotrichia_clavaeformis\|prot_M-clavaeformis_F_contig3058.9953.1 | Myriotrichia_clavaeformis | RP2 | 24.6 | 0.0000000032 |
| Myriotrichia_clavaeformis\|prot_M-clavaeformis_F_contig406.12598.1 | Myriotrichia_clavaeformis | RP2 | 190.8 | 1.10E-60 |
| Myriotrichia_clavaeformis\|prot_M-clavaeformis_F_contig490.14399.1 | Myriotrichia_clavaeformis | RP2 | 187.6 | 9.90E-60 |
| Myriotrichia_clavaeformis\|prot_M-clavaeformis_F_contig1521.3859.1 | Myriotrichia_clavaeformis | TBCC | 181 | 1.30E-57 |
| Myriotrichia_clavaeformis\|prot_M-clavaeformis_F_contig1920.5881.1 | Myriotrichia_clavaeformis | TBCCd1 | 178.4 | 5.90E-57 |
| Nannochloropsis_gaditana\|jgi\|Nangad1\|3894\|rna7282 | Nannochloropsis_gaditana | TBCCd1 | 122.6 | 1.10E-39 |
| Nannochloropsis_oceanica\|Nanoce\|529423\|e_gw1.13.379.1 | Nannochloropsis_oceanica | TBCC | 141.8 | 1.90E-45 |
| Nannochloropsis_oceanica\|Nanoce\|604414\|29_#_184_#_TRIN_DN7881_c4_g1_i4 | Nannochloropsis_oceanica | TBCCd1 | 100.2 | 9.90E-33 |
| Nemacystus_decipiens\|g11173.t1 | Nemacystus_decipiens | RP2 | 183.7 | 1.70E-58 |
| Nemacystus_decipiens\|g11568.t1 | Nemacystus_decipiens | RP2 | 65.9 | 5.00E-22 |
| Nemacystus_decipiens\|g9695.t1 | Nemacystus_decipiens | RP2 | 19.2 | 0.00000014 |
| Nemacystus_decipiens\|g5057.t1 | Nemacystus_decipiens | TBCC | 175 | 9.20E-56 |
| Nemacystus_decipiens\|g5894.t1 | Nemacystus_decipiens | TBCCd1 | 175.9 | 3.60E-56 |
| Nemacystus_decipiens\|g5894.t2 | Nemacystus_decipiens | TBCCd1 | 176 | 3.40E-56 |
| Pelvetia_canaliculata\|prot_P-canaliculata_contig11561.2005.1 | Pelvetia_canaliculata | RP2 | 173.2 | 2.90E-55 |
| Pelvetia_canaliculata\|prot_P-canaliculata_contig66350.20050.1 | Pelvetia_canaliculata | RP2 | 126.9 | 6.20E-41 |
| Pelvetia_canaliculata\|prot_P-canaliculata_contig2275.11091.1 | Pelvetia_canaliculata | TBCC | 84.2 | 1.30E-27 |
| Pelvetia_canaliculata\|prot_P-canaliculata_contig35102.14734.1 | Pelvetia_canaliculata | TBCCd1 | 145 | 1.30E-46 |
| Phaeodactylum_tricornutum\|Phatr3_EG01874.p1 | Phaeodactylum_tricornu | TBCC | 67.3 | 2.20E-22 |

| | | | | |
|---|---|---|---|---|
| | tum | | | |
| Phaeodactylum_tricornutum\|Phatr3_J49350.p1 | Phaeodactylum_tricornutum | TBCCd1 | 119.9 | 7.60E-39 |
| Phaeodactylum_tricornutum\|Phatr3_J50339.p1 | Phaeodactylum_tricornutum | TBCCd1 | 119.9 | 7.50E-39 |
| Phaeosaccion_multiseriatum\|PhaMul_00003853 | Phaeosaccion_multiseriatum | RP2 | 166.8 | 2.80E-53 |
| Phaeosaccion_multiseriatum\|PhaMul_00006777 | Phaeosaccion_multiseriatum | RP2 | 173.2 | 2.90E-55 |
| Phaeosaccion_multiseriatum\|PhaMul_00009910 | Phaeosaccion_multiseriatum | RP2 | 172.6 | 4.70E-55 |
| Phaeosaccion_multiseriatum\|PhaMul_00015031 | Phaeosaccion_multiseriatum | TBCC | 145.7 | 1.10E-46 |
| Phaeosaccion_multiseriatum\|PhaMul_00016041 | Phaeosaccion_multiseriatum | TBCCd1 | 146.3 | 5.20E-47 |
| Phaeothamnion_wetherbeei\|prot_P-wetherbeei_contig10142.1.1 | Phaeothamnion_wetherbeei | RP2 | 160.7 | 2.20E-51 |
| Phaeothamnion_wetherbeei\|prot_P-wetherbeei_contig15161.1.1 | Phaeothamnion_wetherbeei | RP2 | 153.3 | 4.30E-49 |
| Phaeothamnion_wetherbeei\|prot_P-wetherbeei_contig3914.1.1 | Phaeothamnion_wetherbeei | TBCC | 139.4 | 9.70E-45 |
| Phaeothamnion_wetherbeei\|prot_P-wetherbeei_contig13488.1.3 | Phaeothamnion_wetherbeei | TBCCd1 | 163.9 | 1.90E-52 |
| Phytophthora_infestans\|PITG_06877T0 | Phytophthora_infestans | RP2 | 169.3 | 4.70E-54 |
| Phytophthora_infestans\|PITG_13367T0 | Phytophthora_infestans | RP2 | 170.3 | 2.30E-54 |
| Phytophthora_infestans\|PITG_14980T0 | Phytophthora_infestans | RP2 | 171.5 | 1.00E-54 |
| Phytophthora_infestans\|PITG_14614T0 | Phytophthora_infestans | TBCC | 117.8 | 5.00E-38 |
| Phytophthora_infestans\|PITG_01073T0 | Phytophthora_infestans | TBCCd1 | 68.2 | 7.80E-23 |
| Pleurocladia_lacustris\|prot_P-lacustris_contig708.13705.1 | Pleurocladia_lacustris | RP2 | 184.8 | 7.50E-59 |
| Pleurocladia_lacustris\|prot_P-lacustris_contig90.15495.1 | Pleurocladia_lacustris | RP2 | 190.2 | 1.60E-60 |
| Pleurocladia_lacustris\|prot_P-lacustris_contig1788.4797.1 | Pleurocladia_lacustris | TBCC | 139.1 | 1.20E-44 |
| Porterinema_fluviatile\|prot_P-fluviatile_contig12.1461.1 | Porterinema_fluviatile | RP2 | 19 | 0.00000017 |
| Porterinema_fluviatile\|prot_P-fluviatile_contig30.7129.1 | Porterinema_fluviatile | TBCC | 148.2 | 1.90E-47 |
| Porterinema_fluviatile\|prot_P-fluviatile_contig7.13582.1 | Porterinema_fluviatile | TBCCd1 | 178.1 | 7.40E-57 |
| Pylaiella_littoralis\|prot_P-littoralis_Contig224.81.4 | Pylaiella_littoralis | RP2 | 186.7 | 1.90E-59 |
| Pylaiella_littoralis\|prot_P-littoralis_Contig466.18.4 | Pylaiella_littoralis | RP2 | 190.8 | 1.10E-60 |
| Pylaiella_littoralis\|prot_P-littoralis_Contig313.38.5 | Pylaiella_littoralis | TBCC | 74.3 | 1.50E-24 |
| Pylaiella_littoralis\|prot_P-littoralis_Contig593.7.2 | Pylaiella_littoralis | TBCCd1 | 174.3 | 1.10E-55 |
| Pythium_ultimum\|PYU1_T010439 | Pythium_ultimum | RP2 | 176.7 | 2.40E-56 |
| Pythium_ultimum\|PYU1_T014635 | Pythium_ultimum | RP2 | 158.8 | 8.20E-51 |
| Pythium_ultimum\|PYU1_T007905 | Pythium_ultimum | TBCC | 119.5 | 1.50E-38 |
| Pythium_ultimum\|PYU1_T001791 | Pythium_ultimum | TBCCd1 | 57.1 | 2.20E-19 |
| Saccharina_japonica_sp\|SJ17390 | Saccharina_japonica_sp | RP2 | 22.6 | 0.000000013 |
| Saccharina_japonica_sp\|SJ19340 | Saccharina_japonica_sp | RP2 | 190.9 | 1.00E-60 |
| Saccharina_japonica_sp\|SJ12546 | Saccharina_japonica_sp | TBCC | 180 | 2.60E-57 |
| Saccharina_japonica_sp\|SJ20425 | Saccharina_japonica_sp | TBCCd1 | 182.2 | 4.00E-58 |
| Saccharina_latissima\|prot_S-latissima_F_contig1432.3139.1 | Saccharina_latissima | RP2 | 167.1 | 2.30E-53 |
| Saccharina_latissima\|prot_S-latissima_F_contig156.3880.1 | Saccharina_latissima | RP2 | 191.1 | 8.50E-61 |
| Saccharina_latissima\|prot_S-latissima_F_contig1873.5345.1 | Saccharina_latissima | RP2 | 23.4 | 0.0000000072 |
| Saccharina_latissima\|prot_S-latissima_F_contig938.17115.1 | Saccharina_latissima | TBCC | 170.7 | 2.00E-54 |
| Saccorhiza_dermatodea\|prot_S-dermatodea_contig32097.8752.1 | Saccorhiza_dermatodea | RP2 | 37.9 | 2.30E-13 |
| Saccorhiza_dermatodea\|prot_S-dermatodea_contig988.17837.1 | Saccorhiza_dermatodea | RP2 | 177.8 | 1.10E-56 |
| Saccorhiza_dermatodea\|prot_S-dermatodea_contig4605.11795.1 | Saccorhiza_dermatodea | TBCC | 178.7 | 6.80E-57 |
| Saccorhiza_polyschides\|prot_S-polyschides_M_contig10682.604.1 | Saccorhiza_polyschides | RP2 | 18.7 | 0.0000002 |
| Saccorhiza_polyschides\|prot_S-polyschides_M_contig14624.3776.1 | Saccorhiza_polyschides | RP2 | 135.1 | 1.80E-43 |
| Saccorhiza_polyschides\|prot_S-polyschides_M_contig8891.18712.1 | Saccorhiza_polyschides | RP2 | 178.4 | 7.30E-57 |
| Saccorhiza_polyschides\|prot_S-polyschides_M_contig11328.1226.1 | Saccorhiza_polyschides | TBCC | 76.2 | 3.70E-25 |
| Saccorhiza_polyschides\|prot_S-polyschides_M_contig26701.9079.1 | Saccorhiza_polyschides | TBCCd1 | 109.8 | 1.00E-35 |
| Saprolegnia_parasitica\|KDO25737.1 | Saprolegnia_parasitica | RP2 | 175.5 | 5.50E-56 |
| Saprolegnia_parasitica\|KDO32609.1 | Saprolegnia_parasitica | RP2 | 150.9 | 2.30E-48 |
| Saprolegnia_parasitica\|KDO16543.1 | Saprolegnia_parasitica | TBCCd1 | 70.5 | 1.50E-23 |

| | | | | |
|---|---|---|---|---|
| Saprolegnia_parasitica\|KDO27743.1 | Saprolegnia_parasitica | TBCCd1 | 81.5 | 5.90E-27 |
| Sargassum_fusiforme\|SFUSKR1349.13 | Sargassum_fusiforme | RP2 | 29.1 | 1.30E-10 |
| Sargassum_fusiforme\|SFUSKR170.33 | Sargassum_fusiforme | RP2 | 98.8 | 3.40E-32 |
| Sargassum_fusiforme\|SFUSKR92.2 | Sargassum_fusiforme | RP2 | 29.1 | 1.30E-10 |
| Sargassum_fusiforme\|SFUSKR283.8 | Sargassum_fusiforme | TBCC | 165.1 | 1.10E-52 |
| Sargassum_fusiforme\|SFUSKR1145.1 | Sargassum_fusiforme | TBCCd1 | 163.6 | 2.20E-52 |
| Sargassum_thunbergii\|STHUKR1245.6 | Sargassum_thunbergii | RP2 | 49.5 | 6.20E-17 |
| Sargassum_thunbergii\|STHUKR944.4 | Sargassum_thunbergii | RP2 | 28.1 | 2.50E-10 |
| Sargassum_thunbergii\|STHUKR1023.1 | Sargassum_thunbergii | TBCC | 131.2 | 3.50E-42 |
| Sargassum_thunbergii\|STHUKR364.1 | Sargassum_thunbergii | TBCC | 165.7 | 7.30E-53 |
| Schizocladia_ischiensis\|prot_S-ischiensis_contig5.14339.1 | Schizocladia_ischiensis | RP2 | 178 | 9.40E-57 |
| Schizocladia_ischiensis\|prot_S-ischiensis_contig41.12583.1 | Schizocladia_ischiensis | TBCCd1 | 142.8 | 6.50E-46 |
| Scytosiphon_promiscuus_Mr5f\|Mr5f_009813-T1 | Scytosiphon_promiscuus_Mr5f | RP2 | 22.4 | 0.000000015 |
| Scytosiphon_promiscuus_Mr5f\|Mr5f_011283-T1 | Scytosiphon_promiscuus_Mr5f | RP2 | 189.4 | 2.80E-60 |
| Scytosiphon_promiscuus_Mr5f\|Mr5f_012608-T1 | Scytosiphon_promiscuus_Mr5f | RP2 | 192.1 | 4.20E-61 |
| Scytosiphon_promiscuus_Mr5f\|Mr5f_012608-T2 | Scytosiphon_promiscuus_Mr5f | RP2 | 192.1 | 4.00E-61 |
| Scytosiphon_promiscuus_Mr5f\|Mr5f_005052-T1 | Scytosiphon_promiscuus_Mr5f | TBCC | 178.1 | 1.00E-56 |
| Scytosiphon_promiscuus_Mr5f\|Mr5f_003417-T1 | Scytosiphon_promiscuus_Mr5f | TBCCd1 | 177.1 | 1.50E-56 |
| Scytosiphon_promiscuus_Ot16m\|Spro_010468-T1 | Scytosiphon_promiscuus_Ot16m | RP2 | 22.4 | 0.000000015 |
| Scytosiphon_promiscuus_Ot16m\|Spro_011990-T1 | Scytosiphon_promiscuus_Ot16m | RP2 | 189.4 | 2.80E-60 |
| Scytosiphon_promiscuus_Ot16m\|Spro_013346-T1 | Scytosiphon_promiscuus_Ot16m | RP2 | 192.1 | 4.20E-61 |
| Scytosiphon_promiscuus_Ot16m\|Spro_013346-T2 | Scytosiphon_promiscuus_Ot16m | RP2 | 192.1 | 4.00E-61 |
| Scytosiphon_promiscuus_Ot16m\|Spro_005289-T1 | Scytosiphon_promiscuus_Ot16m | TBCC | 177 | 2.30E-56 |
| Scytosiphon_promiscuus_Ot16m\|Spro_003616-T1 | Scytosiphon_promiscuus_Ot16m | TBCCd1 | 176.7 | 2.00E-56 |
| Scytosiphon_shibazakiorum_Os10m\|Sshib_009000-T1 | Scytosiphon_shibazakiorum_Os10m | RP2 | 23.8 | 0.0000000054 |
| Scytosiphon_shibazakiorum_Os10m\|Sshib_010351-T1 | Scytosiphon_shibazakiorum_Os10m | RP2 | 189.4 | 2.80E-60 |
| Scytosiphon_shibazakiorum_Os10m\|Sshib_011586-T1 | Scytosiphon_shibazakiorum_Os10m | RP2 | 192.1 | 4.20E-61 |
| Scytosiphon_shibazakiorum_Os10m\|Sshib_004565-T1 | Scytosiphon_shibazakiorum_Os10m | TBCC | 177.3 | 1.90E-56 |
| Scytosiphon_shibazakiorum_Os10m\|Sshib_003028-T1 | Scytosiphon_shibazakiorum_Os10m | TBCCd1 | 177.1 | 1.50E-56 |
| Sphacelaria_rigidula\|prot_S-rigidula_F_contig14977.1.1 | Sphacelaria_rigidula | RP2 | 178.8 | 5.30E-57 |
| Sphacelaria_rigidula\|prot_S-rigidula_F_contig4218.1.1 | Sphacelaria_rigidula | RP2 | 184.9 | 7.00E-59 |
| Sphacelaria_rigidula\|prot_S-rigidula_F_contig2753.2.1 | Sphacelaria_rigidula | TBCC | 171.7 | 9.60E-55 |
| Sphaerotrichia_firma\|prot_S-firma_F_contig17148.7359.1 | Sphaerotrichia_firma | RP2 | 144 | 3.20E-46 |
| Sphaerotrichia_firma\|prot_S-firma_F_contig3079.14332.1 | Sphaerotrichia_firma | RP2 | 27.7 | 3.50E-10 |
| Sphaerotrichia_firma\|prot_S-firma_F_contig4509.18610.1 | Sphaerotrichia_firma | RP2 | 189.7 | 2.30E-60 |
| Sphaerotrichia_firma\|prot_S-firma_F_contig7499.24211.1 | Sphaerotrichia_firma | RP2 | 153.1 | 5.10E-49 |
| Sphaerotrichia_firma\|prot_S-firma_F_contig4030.17500.1 | Sphaerotrichia_firma | TBCC | 144.6 | 2.50E-46 |
| Sphaerotrichia_firma\|prot_S-firma_F_contig508.19786.1 | Sphaerotrichia_firma | TBCC | 180.9 | 1.40E-57 |
| Sphaerotrichia_firma\|prot_S-firma_F_contig17044.7273.1 | Sphaerotrichia_firma | TBCCd1 | 91.2 | 6.10E-30 |
| Sphaerotrichia_firma\|prot_S-firma_F_contig20217.9407.1 | Sphaerotrichia_firma | TBCCd1 | 21.2 | 0.000000027 |
| Sphaerotrichia_firma\|prot_S-firma_F_contig20345.9482.1 | Sphaerotrichia_firma | TBCCd1 | 52.6 | 5.30E-18 |
| Sphaerotrichia_firma\|prot_S-firma_F_contig23836.11357.1 | Sphaerotrichia_firma | TBCCd1 | 25.1 | 0.0000000018 |
| Sphaerotrichia_firma\|prot_S-firma_F_contig37547.16649.1 | Sphaerotrichia_firma | TBCCd1 | 73.8 | 1.50E-24 |
| Sphaerotrichia_firma\|prot_S-firma_F_contig38170.16840.1 | Sphaerotrichia_firma | TBCCd1 | 54 | 1.90E-18 |
| Thalassiosira_pseudonana\|EED96489 | Thalassiosira_pseudonana | RP2 | 64.2 | 1.70E-21 |
| Thalassiosira_pseudonana\|EED87014 | Thalassiosira_pseudonana | TBCC | 81.3 | 1.00E-26 |
| Thalassiosira_pseudonana\|EED93153 | Thalassiosira_pseudonana | TBCCd1 | 128.7 | 1.50E-41 |

| | | | | |
|---|---|---|---|---|
| Tribonema_minus\|Trimin1\|160141\|e_gw1.12.235.1 | Tribonema_minus | RP2 | 162.6 | 5.60E-52 |
| Tribonema_minus\|Trimin1\|206511\|estExt_Genewise1Plus.C_Ctg_1560025 | Tribonema_minus | TBCC | 116.7 | 1.10E-37 |
| Tribonema_minus\|Trimin1\|312202\|MIX20965_18_43 | Tribonema_minus | TBCCd1 | 139.3 | 7.40E-45 |
| Tribonema_minus\|Trimin1\|320057\|MIX28820_18_43 | Tribonema_minus | TBCCd1 | 139.4 | 7.20E-45 |
| Tribonema_minus\|Trimin1\|347229\|estExt_Genemark1.C_Ctg_90108 | Tribonema_minus | TBCCd1 | 126.9 | 5.10E-41 |
| Undaria_pinnatifida\|Upin_00002726 | Undaria_pinnatifida | RP2 | 183.9 | 1.40E-58 |
| Undaria_pinnatifida\|Upin_00017755 | Undaria_pinnatifida | TBCC | 170.7 | 2.10E-54 |
| Undaria_pinnatifida\|Upin_00008834 | Undaria_pinnatifida | TBCCd1 | 179.7 | 2.40E-57 |

**Table S2. crRNA and primer sequences used for Cas12a genome editing in *Dictyota*.** Top, crRNAs; bottom, primers used to amplify the target regions for genotyping. Protospacer sequences exclude the TTTV protospacer-adjacent motif. In the crRNA names, Ddic refers to Dictyota, ex indicates the targeted exon, the number indicates the position of the protospacer within the exon, and f and r indicate the forward and reverse strand respectively. The guide designations used in the main text and figures are given in the first column.

| Target | Name | Sequence |
|---|---|---|
| APT | Ddic_APT_ex1a_Cas12_cpf1 | GTTATCCTGAACGTGAGCAGG |
| APT | Ddic_APT_ex1b_Cas12_cpf1 | CGCTACTTTCGTTATCCTGAA |
| APT | Ddic_APT_ex2_Cas12_cpf1 | CCTTAGCCGTGAAGGTGTTCG |
| APT | Ddic_APT_ex3_Cas12_cpf1 | ATGCTCGGGGATTCATCCTGG |
| DISTAG | Ddic_DIS_3_56r_Cas12_cpf | GTCGCCCCCCTCCTTCCCGCC |
| DISTAG | Ddic_DIS_3_101f_Cas12_cpf | TGTTCTTGCATCTTCATCCAG |
| DISTAG | Ddic_DIS_5_79f_Cas12_cpf | GGCAGCGGGCCCTCGGCAGCG |
| DISTAG | Ddic_DIS_6_112f_Cas12_cpf | ACGTCGTTGCCCACCGGCGGG |
| DISTAG | Ddic_DIS_6_161r_Cas12_cpf | CTGATCCTCGCTCACCAATAA |
| APT | Ddic_apt2F | CGCCGGCTATAGACACCAAT |
| APT | Ddic_apt2R | ACAGAGCTGTTGCATCTACCC |
| DISTAG | Ddic_DIS_3_F_a | ACACAGTAACGGATCCTGGC |
| DISTAG | Ddic_DIS_3_R_a | AGCTTAACCCGGACATCTCC |

**Table S3. CRISPR/Cas9 mutagenesis of APT i*n Dictyota dichotoma*.** Each row is an independent experiment. Embryos were selected on 2-fluoroadenine (2-FA), and surviving (resistant) embryos were counted as putative apt mutants. Efficiency = resistant / (resistant + dead) embryos. Dead eggs (unfertilised) died before selection and were not included.

| CRISPR date | Guides | 2FA time after fertilisation | Resistance count | Mortality count | % efficiency |
|---|---|---|---|---|---|
| Experiment1 | APT 1A, APT_1b, APT_2, APT_3 | 30 days | 1 | 14 | 6.67% |
| | control | 30 days | no resistance | 20 | all dead |
| Experiment2 | APT_2, APT_3 | 4 days | no resistance | | all dead |
| Experiment3 | APT 1A, APT_1b | 4 days | no resistance | 260 | all dead |
| Experiment4 | APT_2, APT_3 | 4 days | no resistance | | all dead |
| Experiment5 | APT 1A, APT_1b | 5 days | no resistance | | all dead |
| Experiment6 | APT_2, APT_3 | 5 days | no resistance | | all dead |
| Experiment7 | APT 1a&1b | 17 days | 17 | 368 | 4.42% |
| | control | 17 days | no resistance | 107 | all dead |
| Experiment8 | APT 1a&1b | 25 days | 18 | 384 | 4.48% |
| | control | 25 days | no resistance | 127 | all dead |
| Experiment9 | APT 1a&1b | 17 days | 26 | 322 | 7.47% |
| Experiment10 | APT 1a | 34 days | 22 | 61 | 26.51% |

**Table S4. CRISPR/Cas9 mutagenesis of DISTAG in *Dictyota dichotoma*.** Each row is an independent experiment, using either five (DISTAG_ABCDE) or two (DISTAG_AB) sgRNAs. Embryos were scored phenotypically as wild type (wide, light-coloured, flattened blades; putative DIS/DIS), mutant (dark, with a collapsed meristem; putative dis/dis) or showing both phenotypes (putative DIS/dis). Percentages are relative to the total number of scored embryos. Total efficiency = DIS/dis + dis/dis.

| **CRISPR date** | **guides** | **Wild type** | **dark with wild type branch** | **dark and small** | **sum** | **putative DIS/DIS** | **putative DIS/dis** | **putative dis/dis** | **total percentage DIS phenotypes** |
|---|---|---|---|---|---|---|---|---|---|
| Experiment1 | DISTAG_ABCDE | 92 | 7 | 9 | 108 | 85% | 6% | 8% | 14.81% |
| Experiment2 | DISTAG_ABCDE | 24 | 11 | 23 | 58 | 41% | 19% | 40% | 58.62% |
| Experiment3 | DISTAG_ABCDE | 54 | 28 | 55 | 137 | 39% | 20% | 40% | 60.58% |
| Experiment4 | DISTAG_AB | 58 | 25 | 16 | 99 | 59% | 25% | 16% | 41.41% |
| Experiment5 | DISTAG_AB | 34 | 0 | 9 | 43 | 79% | 0% | 21% | 20.93% |

**Table S5. Segregation of the dis phenotype in haploid progeny of edited diploid sporophytes.** Numbers of mutant (dis), normally developing (wt) and mosaic progeny recovered from 3 edited parental sporophytes, shown as counts (top) and as percentages of the progeny scored per release (bottom). Progeny were scored 4 weeks after release. In all parents the two main classes segregated consistent with Mendelian segregation of a single nuclear locus.

| **IN COUNTS** | Rep1 | | | Rep2 | | | Rep3 | | |
|---|---|---|---|---|---|---|---|---|---|
| | *dis* | wt | unclear | *dis* | wt | mosaic | *dis* | wt | unclear |
| *Ddic_dis_C* | 43 | 55 | 2 | 39 | 42 | 3 | 53 | 47 | 0 |
| *Ddic_dis_14* | 53 | 44 | 3 | 53 | 47 | 0 | 49 | 50 | 1 |
| *Ddic_dis_10* | 44 | 55 | 1 | 32 | 28 | 0 | 61 | 38 | 1 |
| **IN PERCENTAGES** | Rep1 | | | Rep2 | | | Rep3 | | |
| | *dis* | wt | unclear | *dis* | wt | branched | *dis* | wt | unclear |
| *Ddic_dis_C* | 43% | 55% | 2% | 46.40% | 50% | 3.50% | 53% | 47% | 0 |
| *Ddic_dis_14* | 53% | 44% | 3% | 53% | 47% | 0% | 49% | 50% | 1% |
| *Ddic_dis_10* | 44% | 55% | 1% | 53.30% | 46.60% | 0% | 61% | 38% | 1% |
